# BET-independent MLV integration is retargeted *in vivo* and selects distinct genomic elements for lymphomagenesis

**DOI:** 10.1101/2022.02.23.481640

**Authors:** Ivan Nombela, Martine Michiels, Dominique Van Looveren, Lukas Marcelis, Sara el Ashkar, Siska Van Belle, Anne Bruggemans, Thomas Tousseyn, Jürg Schwaller, Frauke Christ, Rik Gijsbers, Jan De Rijck, Zeger Debyser

**Author notes:** ZD Corresponding author.

## Abstract

Moloney murine leukemia virus (MLV) infects BALB/c mice and induces T-cell lymphoma in mice. Retroviral integration is mediated by the interaction of the MLV integrase (IN) with members of the bromodomain and extra-terminal motif (BET) protein family (BRD2, BRD3 and BRD4). Introduction of the W390A mutation in MLV IN abolishes BET interaction. Here we compared the replication of W390A MLV and WT MLV in adult BALB/c mice to study the role of BET proteins in replication, integration and tumorigenesis *in vivo*. Comparing WT and W390A MLV infection revealed similar viral loads in blood, thymus and spleen cells. Interestingly, W390A MLV integration was retargeted away from GC-enriched genomic regions. However, both WT MLV and W390A MLV developed T cell lymphoma after a similar latency represented by an enlarged thymus and spleen and multi-organ tumor infiltration. Integration site sequencing from splenic tumor cells revealed clonal expansion in all WT MLV- and W390A MLV-infected mice. However, the integration profile of W390A MLV and WT MLV differed significantly. Integrations were enriched in enhancers and promoters but compared to WT, W390A MLV integrated less frequently into enhancers and more into oncogene bodies, such as *Notch1* and *Ppp1r16b*. We conclude that host factors direct MLV *in vivo* integration site selection. Although, BET proteins target WT MLV integration preferentially towards enhancers and promoters, insertional lymphomagenesis can occur independently from BET, likely due to the intrinsically strong enhancer/promoter of the MLV LTR.

## INTRODUCTION

The integration of the retroviral genome into the host genome is a key step in the replication cycle of retroviruses. Several studies have pointed out that integration site selection of retroviruses is targeted and not a random (1). This event is specific for distinct retroviral families (2). For instance, human immunodeficiency virus type 1 (HIV-1) integration is driven towards active transcriptional units while murine leukemia virus (MLV) integration is directed toward active enhancers and promoters (3–6). The main viral determinant coordinating integration is the integrase (IN). The biochemical reactions of retroviral integration are well understood. After reverse transcription, IN cleaves specific phosphodiester bonds near the viral DNA ends during the 3’ end processing reaction. Then, IN uses the resulting viral DNA 3’-OH groups during strand transfer to cut target DNA. Simultaneously, IN joins each viral DNA end to target DNA 5’-end phosphates. Both reactions proceed via direct transesterification of phosphodiester bonds (7).

The lens epithelium-derived growth factor p75 (LEDGF/p75), is a direct binding partner of HIV-1 IN in and specific for lentivirus integration (8). It tethers the viral pre-integration complex (PIC) to the chromatin (9–12). The tethering mechanism of LEDGF/p75 requires the direct interaction of the integrase binding domain (IBD) at the C-terminus with HIV-1 IN and the recognition of the H3K36m2/3 mark on nucleosomes through the N-terminal PWWP domain (13). Small molecule inhibitors of the interaction between HIV-1 integrase and LEDGF/p75 have been developed and termed LEDGINs (14). Addition of LEDGINs during HIV infection not only reduces integration but also targets residual provirus away from H3K36me2/3 (13, 15). The residual provirus, targeted away from its preferential chromatin landscape, is transcriptionally less active even after reactivation. It appears that retroviruses evolved to adopt epigenetic readers to find preferential integration sites associated with active transcription and/or latency (16).

In the case of MLV, proteins of the bromodomain and extra-terminal domain (BET) family, which include BRD2, BRD3 and BRD4, were identified as binding partners of MLV IN, and proposed to tether the MLV PIC to transcription start sites and enhancers (17–20). BET proteins are composed of two conserved N-terminal bromodomains (BD1 and BD2), an extra-terminal domain (ET) and a C-terminal domain (CTD) which is only present in BRD4 (21). BD1 and BD2 are regions with hydrophobic amino acids able to recognize acetylated H3 and H4 tails (22) and act as chromatin readers. The ET domain associates with a variety of viral and cellular proteins, including transcription factors, chromatin-modifying factors and histone-modifying enzymes (23). The CTD domain is necessary for the recruitment of positive elongation factor (P-TEFb) by the transcriptional complex (24). The MLV IN – BET interaction is mediated through the ET domain of BET proteins and the C-terminal domain of MLV IN (25), and results in the tethering of MLV IN to nucleosomes (26). Amino acids located between positions 390-405 of the MLV IN sequence define a conserved domain in γ-retroviruses involved in the interaction with BRD4 (25). A single substitution of tryptophan for an alanine at position 390 (W390A) in this domain is sufficient to prevent the interaction with BRD2, BRD3 and BRD4 (19) and to shift the integration pattern away from transcriptionally active genomic regions (27). Next to BET proteins, retroviral p12 may also play a role in tethering MLV PICs to mitotic chromosomes to facilitate integration (28).

MLV has been studied as one of the prototype oncogenic animal retroviruses since the 50s, when it was noticed that leukemia could be transmitted to newborn mice by an unknown agent (29). This horizontal transmission occurs primarily through milk ingestion, while transmission by the venereal route is less common (30). MLV induces either lymphoblastic leukemia (30) or lymphoma in mice and an enlarged thymus, spleen or lymph nodes are present in infected adults (31). Since MLV can infect several B or T cells, either leukemia or lymphoma of these cell lineages or their corresponding immature lineages can be induced (32).

Upon integration, MLV can modify mouse gene expression by insertional mutagenesis (33). This phenomenon is characterized by a deregulation of gene expression at the transcriptional or post-transcriptional level (34, 35), and requires viral promoter and enhancer elements located in the U3 region of the long terminal repeats (LTR) of the MLV genome. Therefore, provirus insertion at the start of the gene may enhance transcription of the target gene from the retroviral promoter. Alternatively, viral integration in the central part or close to the 3’ end of a gene may produce a truncated form of the native protein, that can abort its regulatory control. Screening for MLV insertional mutagenesis in mouse models has been instrumental to identify human oncogenes (36–38). Genes identified as targets for retroviral integration that also act as oncogenes are *Notch1, Pim1, Myc, Gfi and Pvt1* among others (39).

Many recent studies on integration site selection of MLV were performed in the context of MLV-based vectors for human gene therapy (40–42). MLV-based vectors have been successfully used in clinical trials for patients with adenosine deaminase deficiency (43). However, serious safety concerns were raised when patients with X-linked severe combined immunodeficiency disease (X-SCID) developed T-cell leukemia after treatment with MLV-based viral vectors by activation of the LMO2 oncogene (44–46). Several generations of viral vectors have been designed in order to increase the biosafety of retroviral therapy. A “second” generation is characterized by a self-inactivating (SIN) vector ensuring a deletion in the 3’ LTR after reverse transcription that abolishes the LTR promoter activity (47). Most recently, a “third” generation of retroviral vectors was proposed. This generation is characterized by a mutation in MLV IN abolishing interaction with BET proteins to achieve an integration profile targeted away from oncogenes (27, 48, 49). Still, our knowledge on the impact of integration site selection on oncogenesis *in vivo* is poor.

In fact, the role of retroviral host factors such as LEDGF/p75 and BET in retroviral pathogenesis in their natural host has not been unambiguously demonstrated. HIV replication in humans in the absence of LEDGF/p75 has not been investigated. Upcoming clinical trials with LEDGINs as HIV treatment or cure may provide this information in the future (50). Loyola *et al*. studied pathogenesis of a C-terminally deleted MLV clone in a tumorigenesis model (51). Unfortunately, the study was somewhat confounded by replication defects of the virus and recombination with endogenous retroviruses.

Here we explored the role of BET-MLV IN interaction in retroviral replication, integration site selection and lymphomagenesis *in vivo*. In contrast to a C-terminal truncation, the site specific W390A mutant replicated to the same levels as wild type MLV. Mice infected with W390A MLV developed T cell lymphoma, to a similar extent as mice infected with wild type (WT) MLV. The W390A mutation targeted integration away from enhancers and promoters, but increased integration in the bodies of known oncogenes. Our observations indicate that loss of BET-MLV IN interaction redirects viral integration sites but does not abrogate MLV-mediated lymphomagenesis. In addition, our work also clarifies the relative contribution of viral integration at enhancers versus promoters or gene bodies in MLV-induced insertional mutagenesis.

## MATERIALS AND METHODS

### Cell culture

Human kidney HEK293T/clone17 was acquired from ATCC (ATCC CRL-11268 293T/17). The NIH/3T3 cell line was a kind gift of Prof. Dr. Johan Van Lint from the Laboratory for Protein Phosphorylation and Proteomics (KU Leuven). Both cell lines were maintained in DMEM – Glutamax I (Gibco, Thermo Fisher Scientific) with 10% fetal bovine serum (FBS) (Gibco) at 37°C and 5% CO2.

### Animal tests

BALB/c mice were acquired from Janvier Labs (Le Genest-Saint-Isle, France). All the experiments and procedures with animals were done according to the European Directive 2010/63/EU for the protection of animals used in scientific purposes. Procedures were also reviewed by the Ethical Commission of Animal tests (Ethische Commissie Dierproeven) from KU Leuven (internal approval number P210/2014, extended P116/2019). An overview of mice used in this study (including a reference to each Figure and experiment) is shown in the Supplementary Table 1.

### MLV constructs

Mutations in the MLV IN of pNCS (52) were done using the SLIM cloning technique (53). Then, the SgrAI/SalI fragment was shuttled into the MLV molecular clone p63.2, which was obtained from Dr. Susan R. Ross (54). p63.2 plasmid is a pBR322 plasmid containing the wild type Moloney MLV provirus clone (55, 56). The C-terminal integrase sequences from the generated MLV clones are shown in Supplementary Figure 1.

### MLV production

Approximately 5.5 ·10^6^ HEK293T cells were plated in a Petri dish. For *ex vivo* experiments, DMEM – Glutamax I (Gibco, Thermo Fisher Scientific) with 10% fetal bovine serum (FBS) (Gibco) and 50 µg/ml Gentamicin (Gibco) were used. For *in vivo* experiments, Opti-MEM I - Glutamax I (Gibco) supplemented with 2% FBS and 50 µg/ml Gentamicin (GIBCO) were used. 25 µg of viral plasmid and 100 µl branched PEI (pH 7.4) (Sigma-Aldrich) were added to the cell culture. After 24 hours post transfection, fresh medium was added. Medium was harvested 48 and 72 hours post transfection and filtered through a 0.45 µM filter (Merck Millipore, Overijse, Belgium). MLV was concentrated by ultrafiltration using Vivaspin MWCP 50 KDa filter tubes (Merck Millipore).

### Semiquantitative PCR and RT-qPCR

RNA extractions were done using the Aurum Total RNA Kit (BioRad) following manufacturer’s instructions. RNA was quantified using the Nano Photometer SP062 (Implen, München, Germany), and 5 µg from each sample was taken for reverse transcription (RT). RT-PCR was done using the High-Capacity cDNA Reverse Transcription kit (Applied Biosystems, ThermoFisher Scientific, Brussels, Belgium).

Semi-quantitative PCR was performed using iProof High-Fidelity kit polymerase (BioRad, Temse, Belgium) following manufacturer’s instructions. PCR was done on the T personal Biometra thermocycler (Westburg). The cycling conditions were 98°C for 45 seconds, 40 cycles at 98°C for 20 seconds, 55°C for 20 seconds, 72°C for 90 seconds ending with 1 cycle at 72°C for 5 minutes. Semi-quantitative PCR products were analyzed by gel electrophoresis using 1% agarose (Cat n° 15510-027, Invitrogen) gel, using the BioRad PowerPac Basic Electrophoresis Power Supply at 140 V for 40 min. The 1 kb GeneRuler (Thermo Fisher Scientific) was used as a ladder.

Quantitative PCR (qPCR) was performed using LightCycler 480 SYBR Green I Master mix (ref.04707516001- Roche) or IQ Supermix (Ref. 1708860, BioRad) for GAPDH gene with a FAM-Tamra probe. Primer concentrations were adjusted to a final concentration of 300 nM. A 2-step qPCR was done using the LightCycler 480 (Roche, Anderlecht, Belgium). The activation cycle was at 95 °C for 10 minutes, followed by 40 cycles at 95 °C for 10 seconds, then 60 °C for 30 seconds. Cooling was done by one cycle at 37 °C during 1 second. Primer sequences used are listed in Table 1.

**Table 1.**
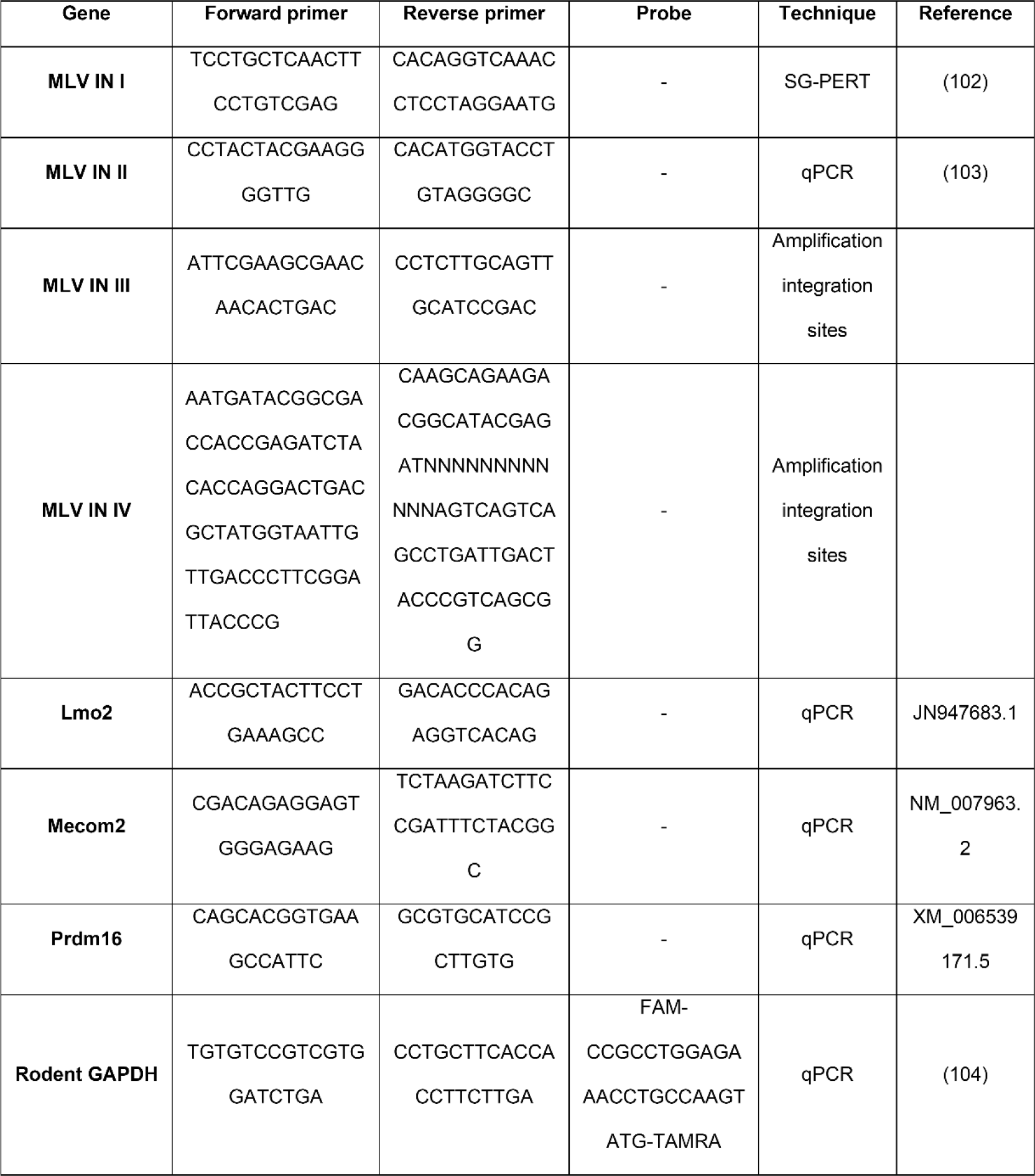
List of primer sequences used for SG-PERT, amplification of integration sites and semi-quantitative PCR.

Table 1. Primer sequences used for SG-PERT, quantitative PCR, semiquantitative PCR and amplification of integration sites.

### MLV titration by SYBR Green product-enhanced reverse transcriptase assay (SG-PERT)

Titration was done as described (57). MLV was produced in NIH/3T3 cells. The supernatant was filtered through a 0.45 µM filter (Millipore). The MLV supernatants and standards were lysed using 2X lysis buffer (0.25% Triton X-100, 50 mM KCl, 100 mM Tris-HCL, 40% glycerol; pH 7.4) while incubated at room temperature for 10 min. After lysis, 1X sample dilution buffer was added to each sample and standard (10X dilution buffer: 50mM (NH4)2SO4, 200 mM KCL, 200 mM Tris-HCl; pH 8.3) with a final dilution of 1/10. Standards were diluted in a dilution series of 1/5. The PCR mix was performed with a final concentration of 0.25 U/µL Fermentas Truestart Hot Start Taq DNA polymerase (Thermo-Fisher) in 100 µL 2X reaction buffer ((5 mM (NH4)2SO4, 20 mM KCl, 20 mM Tris-HCl; pH 8.3), 10 mM MgCl2, 0.2 mg/mL BSA, 1/10000 SYBR green I, 500 µM dNTPs, 4 µM forward primer MLV IN I, 4 µM reverse primer MLV IN I (both listed in Table 1) and 2 µg/ml MS2 RNA. The thermocycler used was the LightCycler 480 (Roche). The cycling conditions were 37°C for 60 min, 95°C for 5 min and 45 cycles at 95°C for 5 sec, 55°C for 5 sec, and 72°C for 15 sec. The final cycle was done at 81°C for 10 seconds.

### Quantification of viral DNA in cells from spleen, thymus and blood

Genomic DNA was extracted from mouse spleen, thymus and blood using the commercial GeneElute Mammalian Genomic DNA Miniprep Kit (Sigma-Aldrich). A 2-step quantitative PCR was performed using MLV IN II primers (listed in Table 1). Rodent GAPDH was used as housekeeping gene using the LightCycler 480 (Roche, Anderlecht, Belgium). The activation cycle was at 95 °C for 10 minutes, followed by 40 cycles at 95 °C for 10 seconds, then 60 °C for 30 seconds. Cooling was done by one cycle at 37 °C during 1 second.

### Quantification of MLV viral titers by co-culture

First, rounded BioCoat^TM^ coverslips 12 mm in diameter (Corning, Fisher Scientific, Merelbeke, Belgium) were added to a 24-well plate (Nunc, Thermo Scientific, Asse, Belgium). Then, approximately 1.5·10^5^ NIH/3T3 cells were seeded in each well. After 24 hours, NIH/3T3 cells were co-cultured with 10^4^ cells from spleen or thymus for 24 hours. Then, NIH/3T3 were rinsed with PBS and they were fixed with 4% paraformaldehyde.

For staining, a primary antibody against MLV protein p12 was obtained from hybridoma alpha CA cells (ATCC CRL-1890). Primary antibody was incubated for 1 hour. Goat anti-mouse biotin (Dako Denmark, Glostrup, Denmark) was used as a secondary antibody and samples were incubated for 20 min, followed by an incubation with streptavidin-HRP (Dako Denmark) for 30 min. Then, a DAB staining was done. Coverslips were mounted using Mowiol®4-88 (Calbiochem, Merck). Infected cells were counted with a Leica DMR microscope (40X magnification).

### Mouse infection

One-day old mice were infected by intraperitoneal injection with 5 or 50 µl of 8·10^4^ RTU/µL MLV, as indicated in each experiment. Moribund mice were sacrificed when the disease reached a late stage, around 12 weeks post-infection by cervical dislocation. Before sacrifice, mice were weighted. Samples of whole blood, thymus and spleen were obtained for histological analysis.

### Blood sampling and cell counting

Blood samples were taken from the submandibular vein by puncture and blood was stored in MAP microtubes coated with 1 mg EDTA (Becton Dickinson, Franklin Lakes, NJ). The blood was diluted 1/10 in PBS and blood cell counting was performed using the hematology analyzer Siemens Advia 2120 (Siemens, Munich, Germany). For further confirmation, blood smears were performed, and blood cells were identified and counted microscopically to corroborate the results from the automatic cell counter.

### Histological sections

Tissues were fixed in 10% neutral buffered formalin for 24 hours and embedded in paraffin. Formalin-fixed and paraffin-embedded sections were cut at 3 micrometer thickness. Automated haematoxylin and eosin (H&E) stains were performed on a Leica ST5010 Autostainer XL (Leica, Wetzlar, Germany) for all samples.

### Integration site sequencing

Integration site sequencing was done as described (58). Briefly, mice were injected with WT MLV or W390A MLV. After 3 and 5 weeks post/infection or at a later stage of the disease (12 – 14 weeks post-infection), mice were sacrificed, and spleens were collected. Genomic DNA was extracted using the GeneElute Mammalian Genomic DNA Miniprep Kit (Sigma). Sonication with the Covaris M220 was used to shear genomic DNA randomly and DNA linkers were ligated to the sheared DNA. Integration sites were amplified by nested PCR using the iProof High-Fidelity kit polymerase (BioRad). Two sets of primers were used in the PCR (sequences are listed in Table 1, indicated as MLV IN III and MLV IN IV), whereby Illumina sequencing adapters were linked to the second set of primers. PCR products were purified by AMPure XP magnetic beads and sequenced by Illumina Miseq (Illumina Inc., San Diego, CA, USA) paired-end 300 cycles by the Leuven Genomics Core (KU Leuven).

### Statistics and software

P values associated with each graph are represented by: *, *P*-value ≤ 0.05; **, *P*-value ≤ 0.01; ***, *P*-value ≤ 0.001; *P*-value ≤ 0.0001. Statistical tests used to analyze significant differences between conditions are indicated in each figure. INSPIIRED software was used to analyze the integration sites (58). Integration site positions were aligned on the mouse genome NCBI37/mm9 to identify insertion in mouse genes using USCS Genome Browser. Genomic size of enhancers in mouse spleen was obtained from EnhancerAtlas 2.0. Genomic size of genes was obtained from the UCSC genes database (assembly July 2007 NCBI37/mm9). Genomic size of mouse promoters was obtained from EPD new promoters database (assembly December 2011 GRC38/mm10). The EnhancerAtlas 2.0 was used to analyze integration into enhancer and promoter features (59). GraphPad Prism 8 was used to make graphs and statistical calculations.

## RESULTS

### Efficient BET-independent *in vitro* replication of the W390A MLV mutant

To investigate the role of BET proteins during MLV replication *in vivo*, we generated two BET-independent Moloney MLV molecular clones: W390A and a C-terminal truncation mutant. Earlier studies revealed that IN W390A is essential for the interaction of IN with BET proteins and as such affects the integration site pattern of MLV (27, 48). In addition, it was shown that this mutation does not hamper the transduction efficiency of MLV-derived viral particles (46). However, its effect on multiple round viral replication was not assessed before. First, a codon optimized sequence duplicating the overlapping sequence at the protein level was inserted in front of the *Env* start codon in molecular clone p63.2. We created WT p63.2 to introduce W390A mutation a single time in MLV IN. Otherwise, introduction of this mutation in the parental p63.2 would have caused selective pressure to revert this mutation due to the presence of the wild type codon in the overlapping Env gene (Supplementary Figure 1A). We also generated ΔC p63.2, carrying a deletion in the C-terminal domain of MLV IN between positions 382 to 408 (Supplementary Figure 2) (60).

HEK 293T cells were transfected with the plasmids p63.2, WT p63.2, W390A p63.2 or ΔC p63.2 to compare the production of viral particles (Supplementary Figure 1B). The reverse transcriptase (RT) activity in the supernatant was measured 48 hours post transfection. Both WT p63.2 and W390A p63.2 were at least as efficient as the parental p63.2 viral clone in the production of viral particles (Supplementary Figure 1C). However, truncation of the C-terminal tail (p63.2 ΔC) resulted in a three-fold decrease in RT activity in comparison with the parental p63.2 clone (Supplementary Figure 1C). Next, NIH/3T3 cells were infected with equal RT units of MLV 63.2, WT 63.2, W390A MLV and ΔC MLV molecular clones (Supplementary Figure 1B). In contrast to the other variants, the ΔC p63.2 clone did not efficiently replicate in NIH 3T3 cells (Supplementary Figure 1D).

Since the C-terminal deletion hampers viral production and replication, this viral clone was omitted from further experiments.

### W390A MLV replicates at wild type levels in BALB/c mice

Considering that WT MLV and W390A MLV can replicate to the same extent *in vitro*, we evaluated their infectivity *in vivo* was evaluated next. Newborn BALB/c mice were injected with 4·10^6^ RTU of WT or W390A MLV, or 50 µL PBS, as indicated in Figure 1A. As a measure of MLV replication, MLV IN RNA expression was analyzed in spleen and thymus samples by RT-qPCR. MLV IN RNA levels in spleen and thymus samples were similar after infection with WT or W390A MLV, although a trend for a lower viral load was seen with the W390A MLV in thymus at 3 weeks post-infection (wpi) (Figures 1B/D). Additionally, *in vivo* infectivity was assessed by co-culture of NIH 3T3 cells with cells of spleen or thymus extracted 3 or 5 wpi from mice infected with WT or W390A MLV. No statistically significant difference was found between the number of infected cells in co-culture with spleen or thymus of either WT- or W390A MLV-infected mice, although a trend for a lower infection from spleen cells infected with the W390A MLV was seen at 3 wpi (Figure 1C/E). To exclude revertants of W390A MLV, we sequenced part of the MLV IN gene in 6 mice infected with parental 63.2 MLV, WT MLV or W390A MLV at 12 weeks post-infection (Supplementary Figure 3). Although some synonymous and non-synonymous base changes were detected, the alanine codon GCA remained intact.

**Figure 1.**
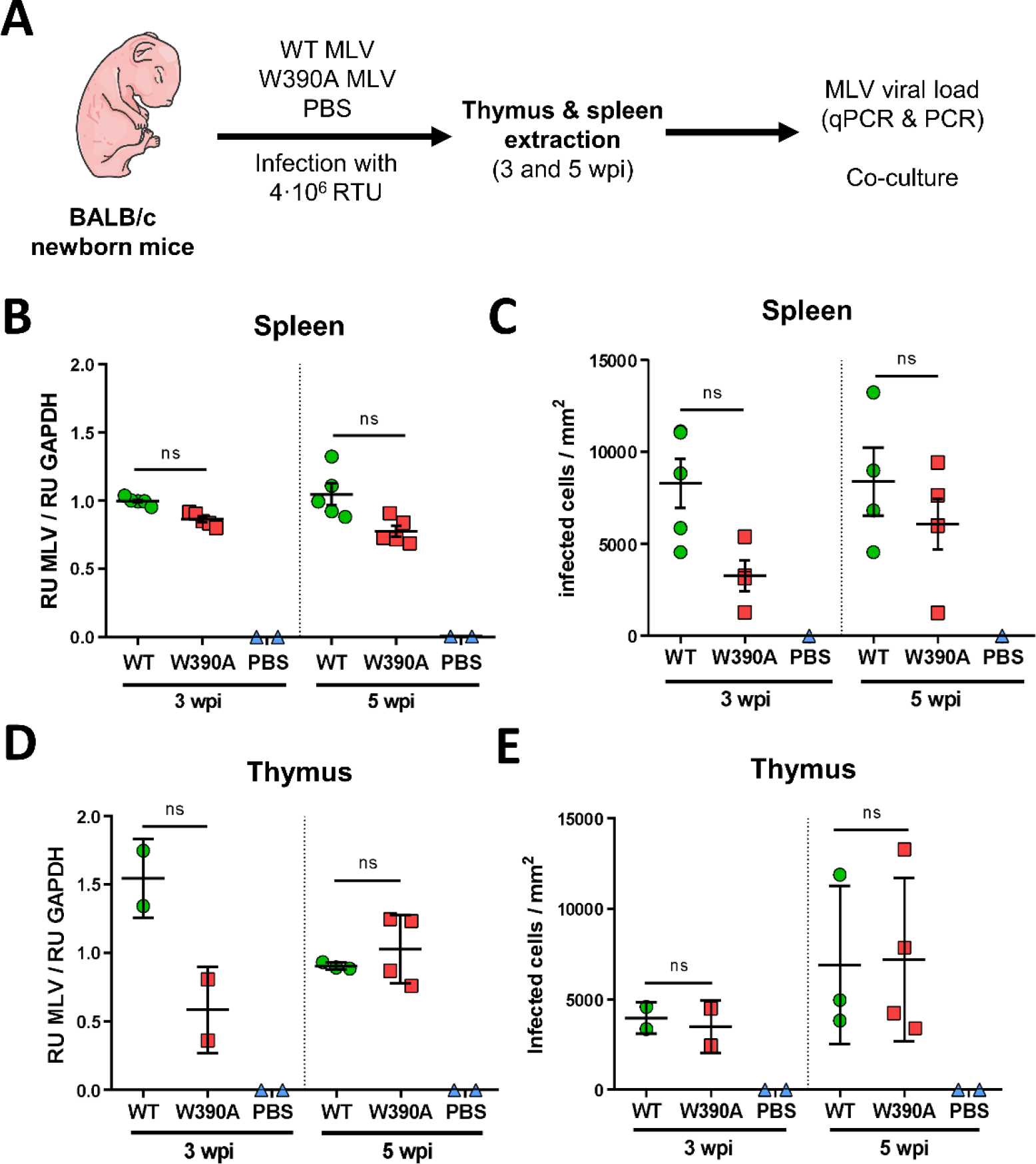
*In vivo* replication of WT MLV and W390A MLV in murine tissue. (**A**) Schematic representation of the work flow to evaluate the *in vivo* infectivity of WT MLV and W390A MLV. Newborn mice were infected by intraperitoneal injection with 4·10^5^ RTU of WT MLV or W390A MLV one day after birth. Blood was drawn between 80 and 90 days post-injection. Extracts of spleen and thymus were made at 3 and 5 weeks post- infection. (**B**) MLV viral load in spleen cells from mice infected with WT MLV (n = 5), W390A (n = 5) MLV or injected with PBS (n = 2) after 3 and 5 weeks post-infection. Viral load was measured by RT-qPCR of MLV IN relative to GAPDH levels. (**C**) Number of infected NIH 3T3 cells after 1 day of co-culture with cells from spleen of mice infected with WT MLV (n = 4), W390A MLV (n = 4) or injected with PBS (n = 2) after 3 and 5 weeks post-infection. (**D**) MLV viral load in thymus cells from mice infected with WT MLV (3 wpi n = 2, 5 wpi n = 3), W390A MLV (3 wpi n = 2, 5 wpi n = 4) or injected with PBS (n = 2) after 3 and 5 weeks post-infection. Viral load was measured by RT-qPCR of MLV IN relative to GAPDH levels. (**E**) Number of infected NIH 3T3 cells after 1 day of co- culture with cells from spleen of mice infected with WT MLV (3 wpi n = 2, 5 wpi n = 3), W390A MLV (3 wpi n = 2, 5 wpi n = 4) or injected with PBS (n = 2) after 3 and 5 weeks post-infection. No statistical significant difference was found using a Kluskal-Wallis test (B-E).

### W390A MLV retargets viral integration

Next, we analyzed integration preferences relative to a set of genomic and epigenetic features. W390A IN is known to shift the integration site preference in cell culture (27, 48). In fact, previous *in vitro* results showed a BET-independent integration profile characterized by lower integration into transcription start sites of RefSeq genes, CpG islands and DNaseI hypersensitive sites (27). Integration sites in spleen samples of mice infected with WT or W390A were analyzed at 3 and 5 weeks post-infection (Figure 2A). Figure 2B summarizes the distribution of integration sites for several genomic features in a 2 kb window. 6000 to 10000 integration sites were obtained for each condition by pooling genomic DNA from spleen cells of three mice infected with either WT or W390A MLV. The presence of W390A significantly shifted the integration away from regions enriched in CpG islands, DNAseI hypersensitive sites, GC-enriched regions and transcription start sites (TSS) (Figure 2C/D), and integration occurred more frequently in positions located within RefSeq genes at both tested times post-infection (Figure 2B). With respect to epigenetic features, W390A MLV integration shifted towards histone marks associated with transcriptionally active and open chromatin such as H3K36me3 (Figure 2D), often related with gene bodies (61). In general, these data demonstrate that the *in vivo* integration site preference of BET-independent MLV in mouse spleen cells differs from that of WT MLV, which is in line with the previous observations in cell culture (27, 48).

**Figure 2.**
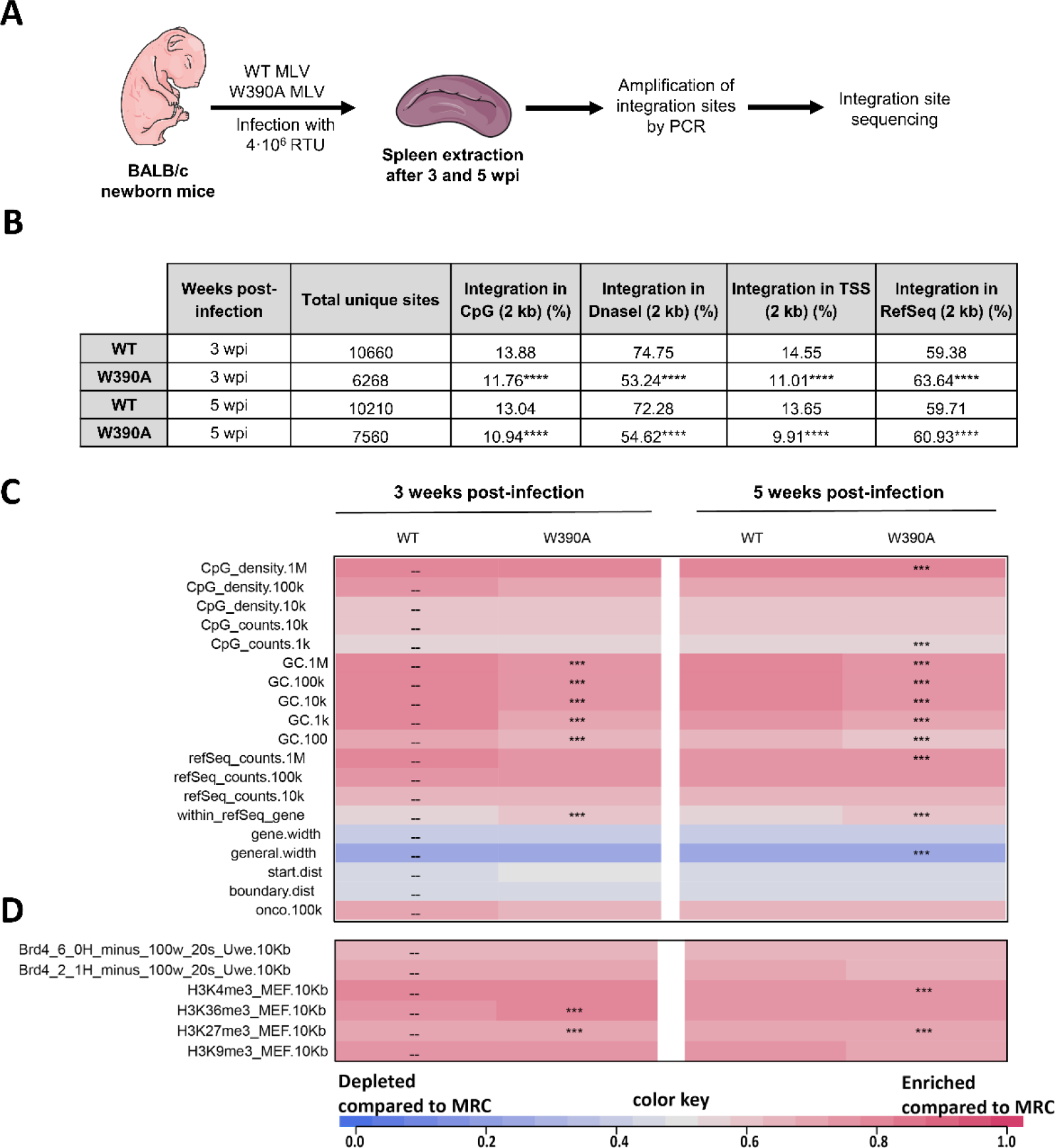
Integration site analysis of WT MLV and W390A MLV in infected mice. (**A**) Schematic representation of the work flow to determine the integration sites and the epigenetic features of WT MLV and W390A MLV integrants *in vivo*. (**B**) Genomic distribution of MLV integration sites obtained from spleen cells of mice infected either with WT MLV or W390A MLV at 3 and 5 wpi. Integration percentages for 2 kb windows around CpG-rich island midpoints, DNAseI hypersensitive sites (DHS), transcription start sites (TSS) and RefSeq genes are given. Statistical analysis was done using a chi square test comparing integrations from WT MLV-infected mice with integrations from W390A MLV-infected mice (*P*-value ≤ 0.0001). Heat maps depicting the relation between integration site frequency and different genomic (**C**) or epigenetic (**D**) features within a 10 kb interval in splenic cells from mice infected either with WT MLV or W390A MLV. The data shown are based on pooled samples from 3 mice infected with either WT MLV or W390A MLV. Features analyzed are shown to the left of the corresponding row of the heat map. Colors indicate whether a particular feature is disfavored (blue) or favored (red) for integration of the respective data sets relative to their computer-generated MRCs, as detailed in the colored scale at the bottom of the panel. Significant departures from WT MLV integration sites in splenic cells are based on a Wald test referred to as χ^2^ distribution.

### Both WT MLV and W390A MLV induce lymphoma in infected mice

Virus-induced hematological malignancies are characterized by increased WBC proliferation (62) and tumor infiltration in several organs and tissues, including lymph nodes, spleen and thymus (63). For this reason, we evaluated the pathology in blood, spleen and thymus from mice infected with either WT or W390A MLV after 12 – 14 weeks post-infection. First, the proliferation of WBCs was assessed as described in Figure 3A. Mice injected with 4·10^5^ RTU MLV (low dose) displayed an increased WBC count (1.46±1.24 x10^4^ for WT MLV and 3.38±4.26 x10^4^ for W390A) compared to mice injected with PBS (0.42 ± 0.12 x10^4^) although this difference was not statistically significant (Figure 3B). Similarly, no statistically significant difference was found between mice infected with WT or W390A MLV. Mice infected with 4·10^6^ RTU MLV (high dose) displayed an increased WBC count (1.22±1.04 x10^4^ for WT MLV and 3.94±4.18 x10^4^ for W390A MLV, respectively). Here the increase in WBC was statistically significant in W390A MLV-infected mice (*P*-value = 0.0044). After low dose infection WT MLV induced an increase in the relative number of neutrophils and monocytes and a decrease in lymphocytes which was not observed with W390A MLV. This phenotype was not seen at high dose (Figure 3B). At high dose, the percentages of monocytes, lymphocytes and neutrophils were similar between mice infected with WT MLV, W390A MLV or PBS (Figure 3C). A summary of the blood cell counts and statistics can be found in Supplementary Table 2.

**Figure 3.**
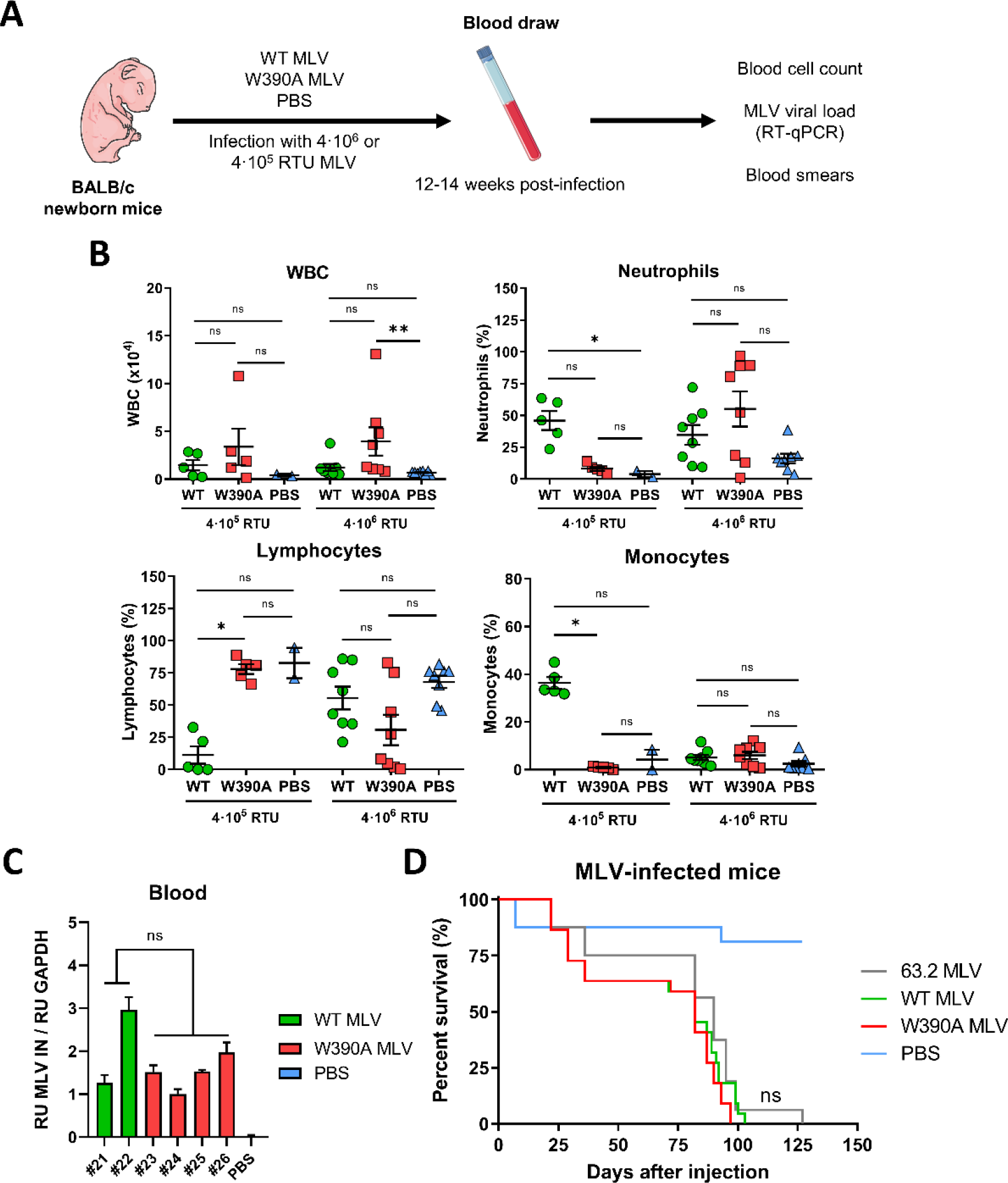
Blood cell counts in WT MLV- and W390A MLV-infected mice. (**A**) Schematic representation of the work flow to evaluate the long-term effect of WT MLV and W390A MLV replication on blood cell numbers. Newborn mice were infected by intraperitoneal injection with 4·10^5^ RTU or 4·10^6^ RTU of either WT MLV or W390A MLV one day after birth. Blood was drawn between 80- and 90-days post-injection. PBS was used as a negative control. (**B**) Number of WBCs in blood samples of mice infected with WT MLV (4·10^5^ RTU n = 5, 4·10^6^ RTU n= 8) or W390A MLV (4·10^5^ RTU n = 5, 4·10^6^ RTU n= 8) or injected with PBS (to compare with 4·10^5^ RTU and 4·10^6^ RTU, n = 2 and 8 respectively). Data correspond to the number of WBCs in 1 µL of blood. Percentages of neutrophils, lymphocytes and monocytes in WBC samples are given for mice infected with WT MLV or W390A MLV. PBS injection was used as negative control. Horizontal lines indicate mean ± SD while dots represent individual values. Statistical significance was analyzed using a Kluskal-Wallis test. (**C**) MLV viral load measured by RT- quantitative PCR of MLV IN gene normalized to GAPDH levels in samples of whole blood from mice infected with WT MLV or W390A MLV taken 21 days post-infection. Each bar represents an individual mouse. Standard deviations were calculated from technical duplicates. (**D**) Survival rates of mice infected with 63.2 MLV, WT MLV or W390A MLV. Newborn mice were infected with 4·10^6^ RTU from each molecular clone diluted in a total volume of 50 µL PBS. Kaplan-Meier plots displaying the survival of mice infected with 63.2 MLV (n = 16), WT MLV (n = 22) or W390A MLV (n = 22). PBS was used as a negative control (n = 16). Each vertical step in the curve indicates one or more deaths. No statistical significant difference was found between 63.2 MLV, WT MLV and W390A MLV using a Gehan-Breslow-Wilcoxon test.

The viral load in whole blood of mice infected with WT (n = 2) or W390A MLV (n = 4) was measured by RT-qPCR detecting IN mRNA at 21 days post-infection. The statistical analysis did not reveal any significant difference (Figure 3C).

We next compared the survival of mice infected with 4·10^6^ RTU WT or W390A MLV. In this experiment, newborn mice were also infected with parental 63.2 MLV. An overview of the different infection experiments is shown in Supplementary Table 3 while Figure 3D shows the pooled data. Mice infected with the distinct MLV viral clones showed similar survival rates (Figure 3D). Although the median survival time was slightly higher for mice infected with MLV 63.2 (90 days) than those infected with either WT MLV or W390A MLV (82 days), no statistically significant difference was found between these MLV clones. A detailed heatmap of the mouse survival including a statistical analysis can be found in Supplementary Table 3. In this figure, mice infected with W390A MLV display a trend towards reduced survival.

### Histopathology of mouse spleen and thymus after MLV infection

Next, we analyzed the pathology of spleen and thymus of mice infected with either WT or W390A MLV (Figure 4A). At 3 and 5 weeks post-infection, prior to the development of lymphoma, spleens from mice infected with WT or W390A were not enlarged (Supplementary Figure 4). In contrast, at late stages of the disease (12 – 14 weeks post-infection), the spleens from mice infected with either WT or W390A MLV were enlarged 20-fold in contrast to spleens from healthy mice injected with PBS (Figures 4B/C). No statistically significant difference was found between WT and W390A MLV.

**Figure 4.**
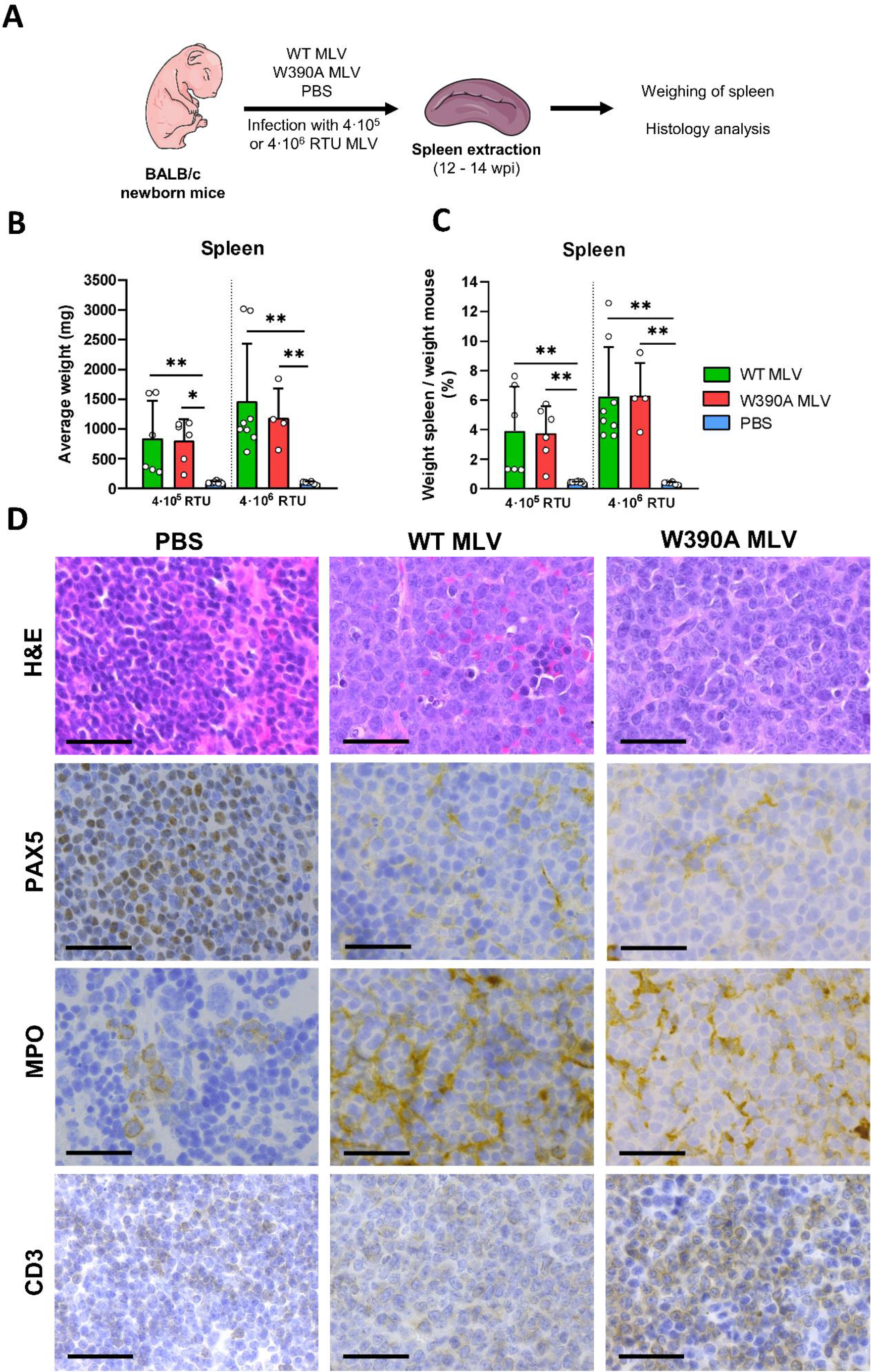
Spleen pathology in WT MLV- and W390A MLV-infected mice. (**A**) Schematic representation of the work flow to analyze the pathology in spleens from WT MLV- and MLV W390A-infected mice. Spleens were sampled between 80 and 90 days post-infection. (**B**) Average weight of the spleen in mice infected with 4·10^6^ RTU of WT MLV (4·10^5^ RTU n = 6; 4·10^6^ RTU n= 9) or W390A MLV (4·10^5^ RTU n = 6, 4·10^6^ RTU n= 4), or injected with PBS (to compare with 4·10^5^ RTU and 4·10^6^ RTU, n = 8 and 6 respectively) as negative controls. Bars show mean ± SD while dots represent individual values. Statistically significant differences were analyzed using a Kluskal-Wallis test. (**C**) Percentage of the weight of the spleen in relation with the body mass of mice infected WT MLV (4·10^5^ RTU n = 6, 4·10^6^ RTU n= 9), W390A MLV (4·10^5^ RTU n = 6, 4·10^6^ RTU n= 4), or injected with PBS (to compare with 4·105 RTU and 4·10^6^ RTU, n = 8 and 6 respectively). Bars show mean ± SD while dots represent individual values. Statistics were done using the Kluskal-Wallis test. (**D**) H&E, PAX-5, MPO and CD3 staining of histological sections of spleen. Mice were infected with 4·10^6^ RTU of WT MLV, W390A MLV or injected with PBS as previously described. Representative images are shown. Black arrows point to mitotic figures. PBS was used as negative control. Scale bar represents 20 µm.

H&E staining of spleen from mice infected with WT MLV or W390A MLV revealed an expansive white pulp with a malignant lymphoid population characterized by large, irregular nuclei and scant cytoplasm (Figure 4D). The irregular nuclei displayed open chromatin and variably prominent nucleoli. Moreover, frequent mitotic figures were present in spleens from mice infected with WT or W390A MLV. Opposite to these findings, spleens from mice injected with PBS displayed normal white and red pulp. In these healthy mice, the white pulp consists of small lymphocytes with rounded nuclei (Figure 4D). We also determined lineage contribution of the tumor cells by staining for PAX-5 (B-cells), myeloperoxidase (MPO) (myeloid cells) and CD3 (T-cells) (64–66). PAX- 5 staining from mice injected with PBS depicted a normal nuclear stain of B cells in a lymphoid follicle (Figure 4D), while MPO staining mainly identified myeloid precursors in the red pulp (Figure 4D). PAX5 and MPO staining of diseased mice showed respectively nuclear and cytoplasmic staining and the malignant population scored negative for either marker. In contrast to MPO and PAX5 markers, CD3 staining of spleen samples from WT MLV-infected mice revealed a relatively weak but extended expression across the sample, while in the spleen samples from mice infected with W390A the staining was stronger. Pathology results are compatible with the development of a T cell lymphoma in mice infected with either WT MLV or W390A.

Tumor infiltration was also observed in lung, liver and kidney tissue from mice infected with WT MLV or W390A MLV (Supplementary Figure 5). Overall, these results demonstrate that the mutant W390A MLV still induces T cell lymphoma. No overt pathological differences were evidenced between BET-dependent (WT) and BET-independent (W390A) MLV although a stronger CD3 staining was evidenced with BET-independent MLV.

### Both WT MLV- and W390A MLV-induced lymphomas display clonal expansion yet a different integration profile

To assess if integration site selection was altered by the W390A mutation, six mice were infected with either WT MLV or MLV W390A and sacrificed at 12 - 14 weeks after infection. In contrast to the previous integration site analysis, at this time point the mice already developed lymphoma due to insertional oncogenesis. The integration profile of both viruses was analyzed as defined in Figure 5A. Briefly, spleen cells from infected mice were isolated and genomic DNA was extracted. Integration sites were determined by ligation-mediated PCR (LM-PCR), followed by high-throughput Illumina sequencing. A summary of the integration sites recovered by this analysis is provided in Supplementary Table 4. Those integration sites are the result of integration site preference and clonal expansion after insertional mutagenesis; we refer to this parameter as “abundance”. An integration site with an abundance ≥ 2 times is indicative of an MLV-infected parental cell that proliferates and generates a daughter cell with the MLV genome integrated in the same position of its genome (67). We refer to these groups of cells as ‘clones’. In Figure 5B, the relative abundance of the clones is displayed. Mice infected with WT MLV presented one dominant clone that represented 10-30% of the total, identified integration sites detected in our analysis. Overall, clones represented 20-50% of the integration sites detected. Other integration sites corresponded to single hits. In the case of W390A MLV, half of the mice showed a different pattern with only few clones present (Figure 5B, mice #8, #9, #10) while the other mice displayed a clonal pattern that resembled that of mice infected with WT MLV (mice #7, #11, #12). No statistical difference was identified between the relative abundance of the three most dominant clones in mice infected with WT MLV or W390A MLV (Figure 5C). However, the distribution of clonal abundance statistically differed between clones generated by WT MLV and W390A MLV integration (Figure 5D). WT MLV integration into intergenic positions generated three groups of clones in the tumors: low abundance (values of 2-3), medium abundance (values between 4 and 6) and high abundance (higher than 10) (Figure 5E). In contrast W390A MLV showed a more polarized profile characterized by a high number of clones with low abundance together with few clones with high abundance regardless of integration in intergenic positions, gene bodies or promoters (Figure 5E). These results indicate that W390A MLV still induces clonal proliferation of lymphoma cells.

**Figure 5.**
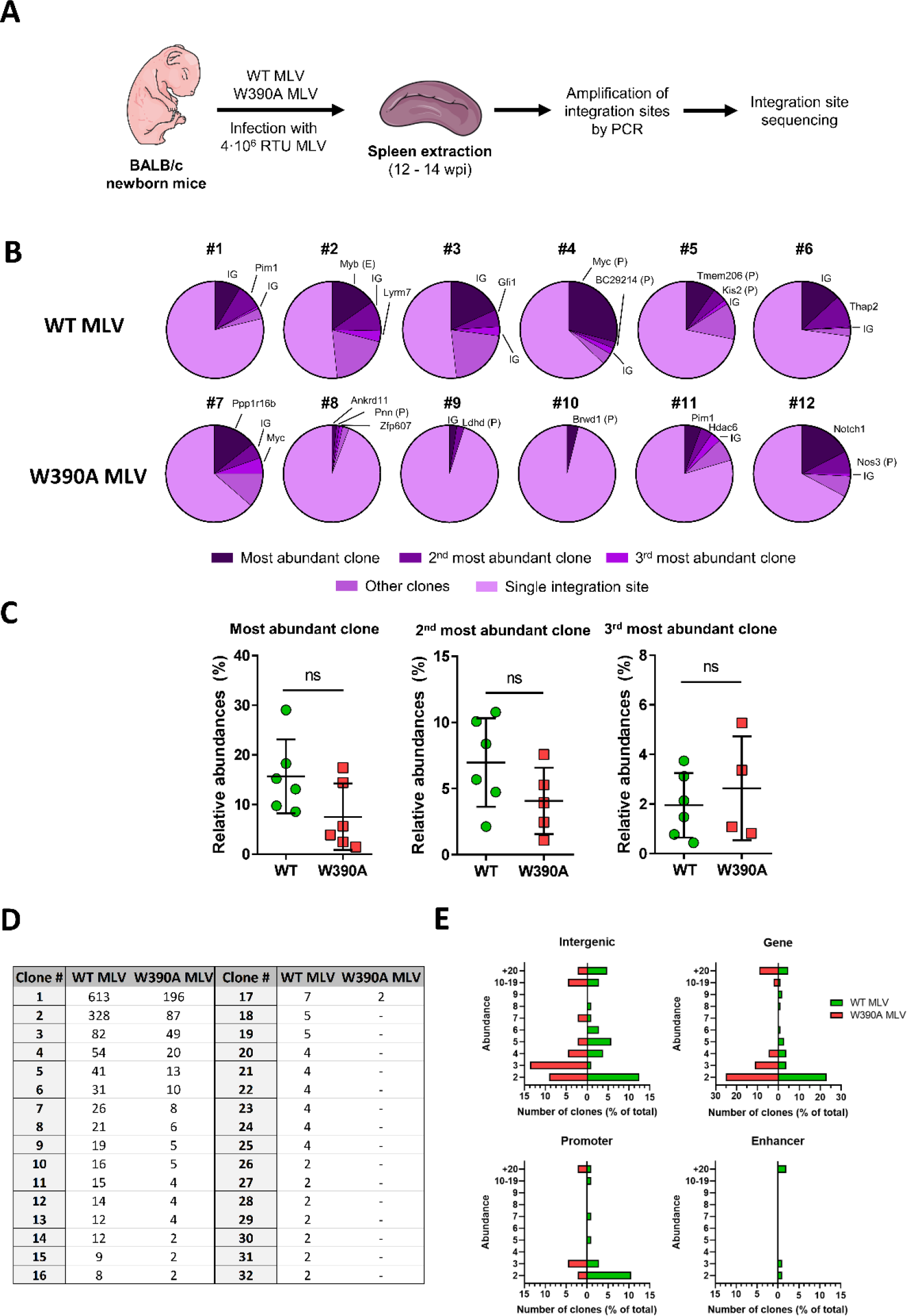
Clonal expansion of tumor cells in spleen induced by WT MLV or W390A MLV. (**A**) Schematic representation of the work flow to determine the integration site selection of WT MLV and MLV W90A in mouse tumor cells after 12 - 14 weeks post- infection. (**B**) Pie charts representing the relative abundance of clones and single integration sites. Each pie chart corresponds to a single mouse infected with WT MLV or W390 MLV. Intergenic site (IG), gene, enhancer (E) or promoter (P) names indicate MLV integration site of the three most dominant clones in each mouse. (**C**) Relative abundance of the top three integration sites in mice infected with WT MLV or W390A MLV. No statistically significant difference was found using a Mann-Whitney test. (**D**) Distribution of clonal abundance in tumors from mice infected with WT MLV or W390A. Clones are ordered from the most dominant (clone #1) to less dominant (clone #32). Each row represents the total abundance of single clone, ordered from more dominant to less dominant. Total abundance was calculated as the sum of such clone from six mice infected with WT MLV (mice labelled from #1 to #6) or W390A MLV (from #7 to #12) respectively. Statistical analysis was done using a chi square test comparing clonal distribution from WT MLV-infected mice with W390A MLV-infected mice (*P*-value ≤ 0.0001 (**E**) Relative number of clones with a specific abundance in tumors induced by WT MLV and W390A MLV integration into intergenic positions, gene bodies, promoters and enhancers.

Next, we investigated the integration sites having a clonal origin in more depth. From the chromosome positions provided in Supplementary Table 4, we identified genes targeted by MLV integration using an alignment to the NCBI37/mm9 mouse genome. WT MLV integration is characterized by a higher number of clonal integration sites (1365 clones of WT MLV versus 284 clones for W390A MLV), (Figures 6A). Both WT MLV and W390A MLV proviruses integrated in *Myc*, *Notch1*, *Pim1*, *Ppp1r16b* and *Thap2* oncogenes. In comparison with WT MLV, W390A MLV integrated less in intergenic positions, but more frequently in the gene bodies of *Notch1* and *Ppp1r16b*. The chromosomal analysis of the MLV integration profile revealed that WT MLV preferentially integrated in chromosomes 5, 7, 10 and 12 while W390A MLV preferred chromosomes 2, 15, 17 and X (Supplementary Figure 6A). However, neither WT nor W390A MLV integration into the host genome was driven by gene density (Supplementary Figure 6B). In summary, these results reveal that lymphoma cells induced by W390A MLV display a significantly different integration profile than lymphomas induced by WT MLV.

**Figure 6.**
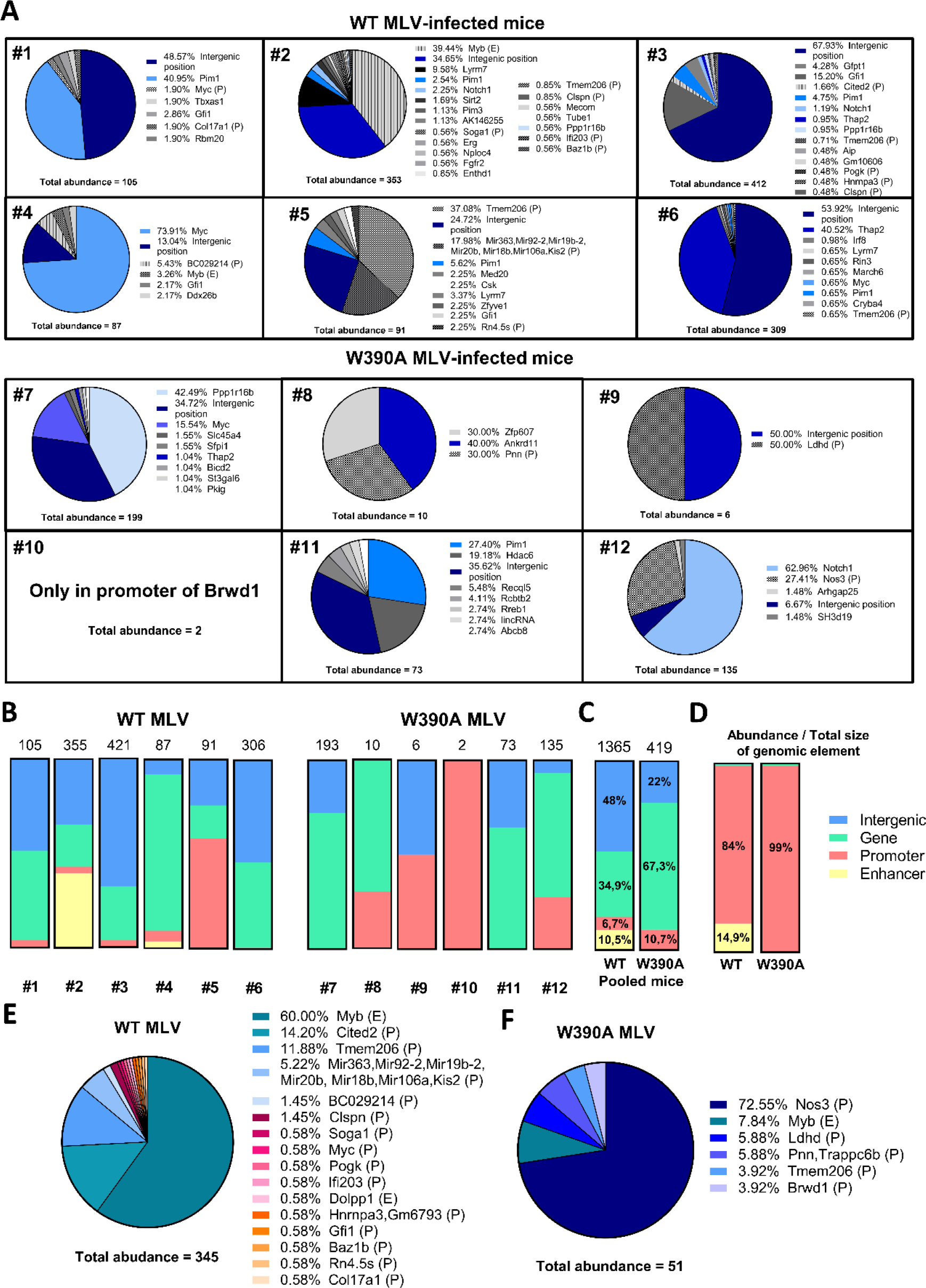
Distribution of MLV abundance in clones generated by WT MLV or W390A MLV insertion. We refer to MLV abundance as the relative frequency of integration sites in expanded clones; this abundance is the result of insertion site preference and clonal expansion induced by the integrated provirus. (**A**) Pie charts depicting individual gene insertional profile of splenic cells with clonal expansion from mice infected with 4⋅10^6^ RTU of either WT MLV or W390A MLV. Total abundance detected by NGS analysis is shown below each pie chart for both MLV constructs. The percentage before each gene name indicates the relative abundance of this gene. Genes with insertions of both WT MLV and W390A MLV are colored in a blue scale. (**B**) MLV abundance into intergenic positions (blue), gene bodies (green), promoters (red) and enhancers (yellow) in WT MLV- and W390A MLV-infected mice. The numeric mouse label is indicated below each bar. Total abundance of clonal integration sites in each mouse is indicated on top of the bar. (**C**) Pooled MLV abundance from intergenic positions (blue), gene bodies (green), promoters (red) and enhancers (yellow) in WT MLV- (n = 6) and W390A MLV-infected mice (n = 6). Total abundance of pooled clonal integration sites from mice infected with either WT MLV or W390A MLV is indicated on top of the bar. Distribution of integration sites between WT MLV and W390A MLV was analyzed using a chi-squared test. The test reported a statistically significant difference between both distribution (*P*-value ≤ 0.0001). (**D**) MLV abundance from intergenic positions (blue), gene bodies (green), promoters (red) and enhancers (yellow) normalized by the total genomic size of each genomic feature. Data was normalized by dividing the percentage of integration site preference into each genomic feature by its own estimated total genomic size: intergenic positions (58.97%), genes (40.54%), promoters (0.05%) and enhancers (0.44%). Data represent six mice infected with in WT MLV or W390A MLV. (**E-F**) Abundance in enhancer (E) or promoter (P) elements in spleen cells from mice infected with 4⋅10^6^ RTU of either WT MLV (n = 6) or W390A MLV (n = 6) isolated after 12 - 14 weeks post- infection. Total abundance of integration at E and P sites is shown below each pie chart for both MLV constructs. The percentage before each gene name indicates the relative abundance of this gene. Integration into an enhancer or promoter element is indicated with an (E) or (P).

### W390A MLV is targeted away from promoters and enhancers *in vivo*

We further analyzed if these integration sites correspond to enhancer or promoter elements using EnhancerAtlas 2.0. Figures 6B and 6C show MLV integration site preference into gene bodies, intergenic positions, enhancers and promoters after clonal expansion. In comparison, WT MLV-induced lymphoma clones displayed a relatively higher abundance of integration into enhancers (as shown for individual mouse #2 and mouse #4) and intergenic positions (on pooled data, 10.5% and 48%% of the total abundance, respectively), while W390A MLV induced clones displayed a higher abundance in promoters (as shown in individual mouse #9 and #10) or gene bodies (mouse #7, #8, #11 and #12) (on average, 10.7% and 67.3% of the total abundance, respectively). Strikingly, integration into promoters and enhancers was particularly enriched, considering their small size in the genome. Normalizing to genomic size showed that WT MLV has a 84% abundance into promoters and a 14.95% abundance into enhancers whereas W390A MLV shows a 99% abundance in promoters (Figure 6D) (Table 2). Next, we pooled all the abundance values into enhancers and promoters from the mice infected with either WT MLV or W390A MLV in the same pie chart (Figure 6E/F). WT MLV showed preferentially clonal expansion from integration in the *Myb* enhancer (60% of the total integration sites into enhancer or promoter elements) and the *Cited2* (14.2%) and *Tmem206* (11.88%) promoters (Figure 6A). Integration into the *Myb* enhancer and the *Tmem206* promoter were also detected in W390A MLV-infected cells, but at a lower frequency (7.84% and 3.92%, respectively) (*P*-Value < 0.0001, χ^2^ test comparing distributions).

**Table 2.** Enrichment of integration in enhancers and promoters for WT MLV and W390A MLV.

|  |  | % Integration / Genomic size |  |
| --- | --- | --- | --- |
|  |  | Promoter | Enhancer |
| High dose | WT MLV | 83.99 | 14.95 |
|  | W390A MLV | 99.05 | 0 |
| Low dose | WT MLV | 49.85 | 48.79 |
|  | W390A MLV | 76.11 | 17.96 |

The analysis of the MLV integration profile from the individual mice revealed that each mouse has a unique integration profile driving the development of the lymphoma (Supplementary Figure 7). In the case of W390A MLV, mouse #7, #11 and #12 had a dominant clone in the tumor with integrations into *Ppp1r16b*, *Pim1* or *Notch1* oncogenes, respectively. In the case of WT MLV, mouse #2 displayed a dominant integration at the *Myb* enhancer while mouse #5 showed integrations into the *Tmem206* promoter. The lymphoma developed by mouse #4 was strongly driven by integration into the *Myc* oncogene.

To determine the potential impact of viral titer, we also analyzed integration site selection of WT MLV and W390A MLV in two groups of three mice infected with a lower dose (5·10^5^ RTU) (Supplementary Figure 8A). The results corroborate the findings in mice infected with a higher dose. Mice infected with W390A MLV at a lower dose displayed lower abundance into enhancers and promoters (Supplementary Figure 8B). When pooling the data from three mice, 3.1% of of the total abundance was found related to enhancers, while 26.7% of the total abundance was found in promoters from WT MLV-infected mice (Supplementary Figure 8C). In contrast, mice infected with W390A MLV displayed only 1.3% of total abundance in promoters and 2.7% in enhancers (Supplementary Figure 8C). Similar to mice infected with a higher dose of MLV, integration and clonal expansion from enhancers and promoters were enriched (for WT MLV, 49.9% in promoters and 48.9% in enhancers, respectively; for W390A MLV, 76.1% in promoters and 18% in enhancers) (Supplementary Figure 8D) (Table 2). All mice displayed a polyclonal integration profile, except for mouse #29 that displayed a oligoclonal integration profile (Supplementary Figure 8E). Integration into the *Myb* enhancer was found in WT-MLV infected mice (mouse #29 and #30). In mice infected with W390A MLV, integration into *Notch1* and *Pvt1* oncogene bodies was evidenced (Supplementary Figure 8F).

Finally, we analyzed if MLV insertional mutagenesis resulted in truncated oncogenes using NGS analysis of WT - or W390A MLV-infected mice. Integrations of both WT or W390A MLV into *Thap2*, *Pim1* and *Notch1* occurred towards the 3’-end of the gene body (Supplementary Figure 9A). However, both WT and W390A MLV integrated in the central part of the *Ppp1r16b* gene. The integration preference of WT MLV and W390A differed with respect to the *Myc* gene; whereas WT MLV integrated near the 3’-end, W390A MLV integrated near the 5’-end. Previous studies on the mechanism of insertional mutagenesis reported that MLV insertion near gene bodies can increase gene expression due to intrinsic enhancer and promoter elements within the long terminal repeats (LTRs) of MLV (6, 68). Common targets for MLV integration are regions nearby *Lmo2*, *Mecom2* and *Prdm16* genes in human (69) or mouse (68). Intragenic insertion in these genes was not identified in our integration site analysis, but we analyzed the expression of these genes using RT-qPCR in spleen cells from a different set of MLV-infected mice isolated at 12 to 14 weeks post-infection by qPCR. The results show that MLV 63.2, WT MLV and W390A MLV do effectively increase the expression of *Mecom2* and *Prdm16* (Supplementary Figure 9B). However, 63.2 MLV and WT MLV were able to induce a stronger increase in *Mecom2* and *Prdm16* expression than W390A MLV.

Although most WT MLV integrations took place into intergenic positions and gene bodies, there was a relative enrichment for enhancers and promoters that only represent 0.49% of the mouse genome. The proportion of integration into enhancers and promoters in mice infected with WT MLV was 17.2% with the high dose and 29.8% with the low dose. BET-independent W390A MLV IN integrated away from enhancers in tumorigenic cells. At a high dose only 10.7% of W390A MLV integrated in enhancers/promoters and with a low dose only 4%.

## DISCUSSION

### BET proteins are not required for MLV *in vivo* integration and replication

Our work unequivocally demonstrates that the interaction between BET proteins and MLV IN is not essential for MLV integration, replication and lymphomagenesis *in vivo*. We reported previously that the W390A mutation abrogates total interaction with BET proteins using an AlphaScreen assay, to a similar level as the MLV IN1-381 construct that lacks the C-terminal tail of the MLV IN (19). In the present study, we demonstrate that W390A MLV can replicate in *in vivo* conditions, which strongly suggest that additional host or viral factors may be involved during integration. While the potential alternative co-factor remains unknown, the finding is in line with our previous observations for HIV integration. We and others reported that HRP-2 (HDGF2) can substitute for LEDGF/p75 in LEDGF/p75 knock-out cells (70, 71).

Our data show that the BET-independent MLV W390A mutant integrates and replicates in mice to a similar extent as the WT virus. This suggests that either MLV IN directly interacts with chromatin or uses alternative host factors or mechanisms. A likely candidate is p12, a cleavage product of MLV Gag, that has been implicated in the early steps of the replication cycle of MLV (72, 73) as well as in virion production (74). The MLV protein p12 can tether the MLV pre-integration complex to host chromosomes (75, 76) by phosphorylation of serine 61, the key event to tether the MLV pre-integration complex to chromatin (49, 77). Yueh and Goff confirmed that lack of p12 phosphorylation of the serine amino acids at positions 61 and 78 result in impaired formation of LTR circles (52), but these mutants could still release normal levels of mature virions, suggesting that p12 may not be essential for the replication of MLV. The relative roles of viral p12 and cellular BET proteins in tethering and targeting integration await further investigation. Possibly tethering by p12 of MLV PICs to mitotic chromosomes is an essential feature while BET-mediated targeting is an optional feature, that still may have evolutionary advantages, since BET proteins remain bound to mitotic chromosomes and are associated with early gene transcription after completion of mitosis (78).

### The MLV-BET interaction is not the essential driver of tumorigenesis

BALB/c mice infected with either WT MLV or W390A developed the same disease phenotype. Upon evaluation, we identified an enlargement of the thymus and spleen, both typical for lymphoma (79). Histopathology identified the same type of malignant population by H&E staining in both WT and W390A MLV-infected mice. These malignant cells are transcriptionally and mitotically more active compared to cells from mice injected with PBS. MPO and PAX-5 staining revealed a lack of B cells, granulocytes and monocytes in the spleen lymphoma. Histopathology and marker analysis revealed strong CD3 staining in mice infected with WT or W390A MLV in contrast with mice injected with PBS, indicating that those mice developed a lymphoma of T cells precursors (80). Moreover, the lymphomas were characterized by polyclonal expansion. Lymphomas induced by retroviruses often display a polyclonal phenotype (81), due to integration into multiple loci in the same cell, that, eventually, lead to oncogenesis by disruption of gene networks that control proliferation (82). Apparently, 1 of 6 mice (mouse #10) infected with W390A MLV displayed a monoclonal lymphoma.

### The lack of BET-MLV IN interaction still yields lymphomas through insertional mutagenesis in oncogene bodies

In the context of gene transcription, BRD4 plays an important role at super-enhancers (SEs) (83). SEs have been defined as clusters of enhancers that are occupied by a high density of multiple transcription factors (84). In this sense, BRD4 and MED1 can act as coactivators of the SEs, and their presence defines the presence of a SE (85). In cancer cells, the acquired cancerous phenotype relies on the abnormal transcription promoted by super-enhancers (83). Although BRD4 may acquire this function at later stages when the lymphoma develops in MLV-infected mice, the tethering mechanism whereby MLV IN binds to BRD4 may point to a role during the retroviral integration step. We show that MLV preferentially integrates into promoters and enhancers under *in vivo* conditions, which is in line with results previously published by other groups in cell lines and primary cells (5, 6, 86). For WT MLV, the integration into enhancers was strongly enriched, while W390A MLV integration was enriched in promoters and gene bodies. Normalization of the relative abundance in Figure 6C for genomic size resulted in an integration preference higher than 98% (WT MLV = 98.9%; W390A MLV = 99%). Whereas WT MLV preferentially integrates in promoters and enhancers, BET-independent MLV preferentially integrates in promoters. The role of BDR4 is to target integration into enhancers, known to interact with BRD4.

We found that the integration of WT MLV into the enhancer of *Myb* (87), and the promoters of *Tmem206* and *Cited2*, induced a stronger clonal expansion. Both Tmem206 and Cited2 were recently described as oncogenes involved in lymphoma (88, 89) in human cells. Overexpression of *Myb* has been detected in both acute myeloid leukemia and non-Hodgkin lymphoma (90, 91), while *Tmem206* and *Cited2* overexpression were detected in colon and prostate cancer, respectively. The molecular mechanism implicated in oncogenesis driven by *Tmem206* is the activation of the AKT signaling pathway (92). Moreover, both the parental 63.2 MLV and WT MLV enhanced expression of *Mecom2* and *Prdm16* (Supplementary Figure 8) while the increase in expression induced by W390A MLV was 2-fold lower, compatible with a lower integration frequency of W390A MLV near enhancers and/or promoter elements. The W390A mutation into MLV IN significantly retargeted viral integration away from enhancers towards oncogenes bodies, such as *Notch1* and *Ppp1r16b*. *Notch1* has been widely involved as one of the driver genes in T-ALL (93–95), while *Ppp1r16b* can promote aggressive lymphomas in mice (96). The truncation of *Notch1* can generate a lymphoma by either loss of the PEST domain located in the 3’-end of the *Notch1* gene, which leads to an enhanced protein half-life and therefore, longer activity as transcription factor in the nucleus (97). Insertional mutagenesis of *Notch1* can also result in deletions in the 5’-end that activate intragenic promoters driving the expression of truncated transcripts that lack the negative regulatory region (NRR) (98). In the case of *Ppp1r16b*, a previous study reported a modest increase in the expression of *Ppp1r16b* in several MLV-induced lymphomas (96). Such increase was more relevant when MLV insertion happened in the *Ppp1r16b* locus.

As such, our results indicate that tumors induced by WT MLV are frequently (44.4%, 4 out 9 mice, corresponding to mouse #2, #5, #27, #28) caused by an overexpression of oncogenes (such as *Myb*) resulting from an integration in an enhancer or promoter near such oncogene, while the tumors induced by W390A MLV are generated frequently (77.7%, 7 out of 9, corresponding to mouse #7-8, #11-12, #30-32) after insertional mutagenesis in an oncogene body (Figure 7).

**Figure 7.**
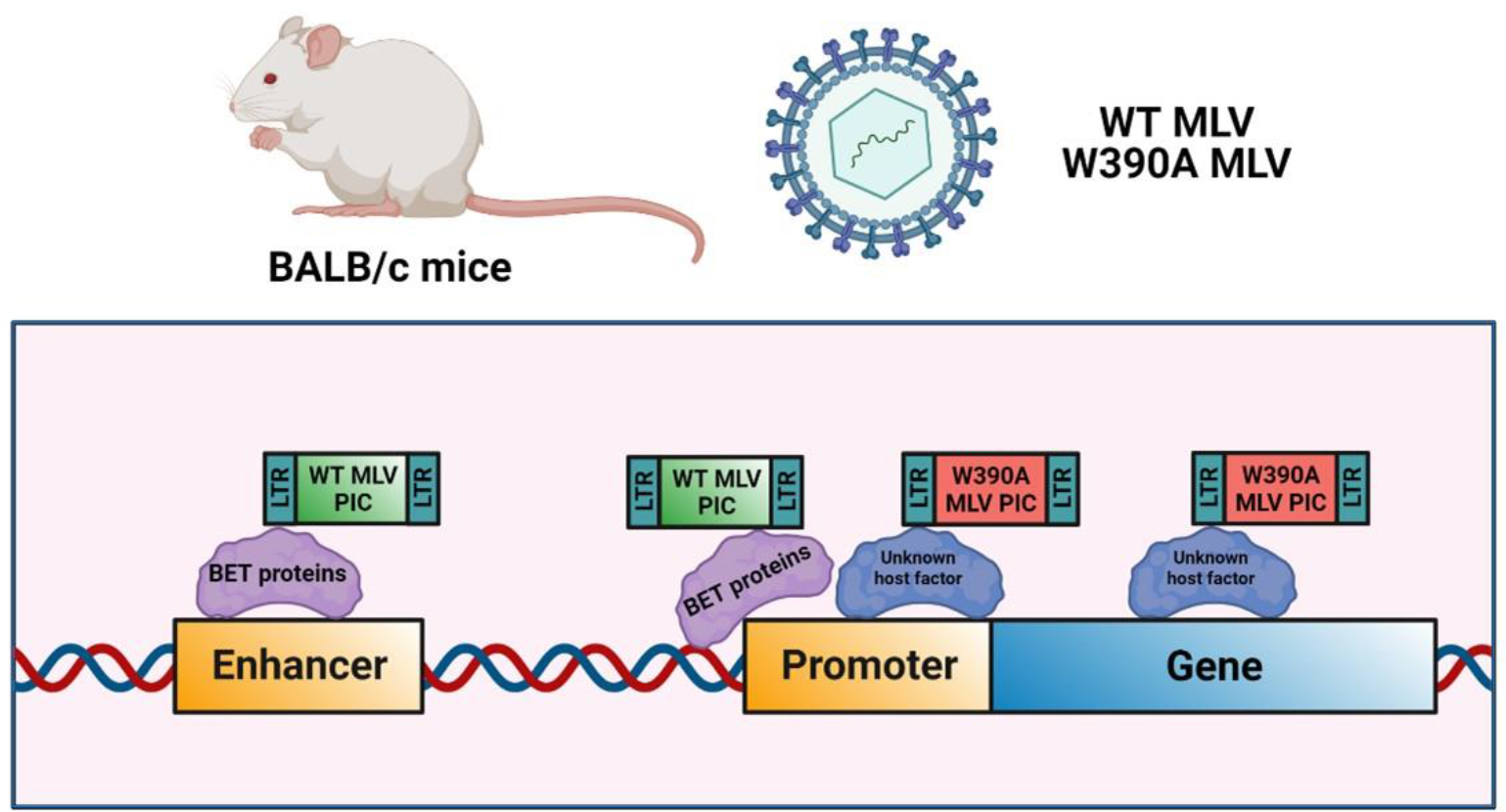
Model for MLV integration *in vivo*. The MLV pre-integration complex (PIC) is targeted towards enhancers and promoters by the BET proteins, while lack of interaction between MLV PIC and BET proteins retargets the integration towards promoters and gene bodies. The strong MLV LTR enhancer/promoter can drive oncogenesis even after retargeting.

### Importance of the C-terminal tail of MLV IN in viral replication

The role of MLV IN-BET protein interaction in an *in vivo* context was previously studied by the group of M. Roth (51). In contrast with our current study with the W390A mutant, distinct BET-independent MLV constructs were used. First, carrying a truncation of 23 amino acids from the C-terminal tail of MLV IN (MLV IN XN) (99), deficient for BRD4 interaction was generated. Next, the MLV IN Terminal Peptide^-^ construct was made including stop codons in the non-*Env* reading frames to prevent restoration of WT IN. The tumorigenesis mouse model MYC/Runx2 was used by the Roth group (51), that is characterized by the spontaneous development of lymphoma (100). As such it does not reflect primary insertional lymphomagenesis upon MLV infection. We used healthy BALB/c mice instead that only develop tumors upon MLV infection. Both MLV IN XN and MLV IN TP^-^ constructs showed a 10-fold decrease in *in vitro* infectivity which corresponds more or less with the 25-fold decrease in infectivity after 24 hours post-infection in NIH 3T3 cells seen with ΔC p63.2 MLV, lacking 27 C-terminal amino acids (Supplementary Figure 1D). While W390A MLV replicated to WT levels in our mouse model, no evaluation of the infectivity under *in vivo* conditions was performed by Loyola and colleagues but it can be deduced that replication of MLV with IN c-terminal deletions was hampered.

Whereas W390A MLV induced lymphomagenesis trended to reduce mouse survival, the TP-deleted MLV clones from Loyola and colleagues increased survival of *MYC/Runx2* mice with 15 days (51). This result may however reflect reduced virus replication *in vivo*. In the latter study, recombination with endogenous retrovirus *Pmv20* was able to revert the expression of the terminal peptide, confounding the analysis (51). We used BALB/c mice, characterized for the presence of polytropic and xenotropic endogenous retroviruses (101). Although mice infected with W390A MLV displayed a higher number of single mutations in MLV IN, suggesting selective pressure, no revertants due to mutagenesis or recombination were observed (Supplementary Figure 2). As for the integration site analysis, both studies report less integration into CpG islands and TSS with BET-independent virus. In conclusion, both studies report BET-independent MLV integration in mice. In both studies BET-independent MLV is retargeted but still associated with lymphoma. In the previous study poor replication capacity and recombination, hampered the analysis of the role of BET interaction in lymphomagenesis while our set up allowed a more detailed pathological and genomic analysis.

### Genotoxicity of BET-independent MLV vectors

One important application of MLV integration studies is the development of safer MLV-based vectors for gene therapy. Previously, it was shown that the safety of SIN MLV vectors varies in different tissues (71). In this study, we have used the W390A mutation in MLV IN to get a better understanding of the relevance of BET proteins in the development of lymphoma. Overall, our results indicate that prevention of the MLV IN – BET interaction was sufficient to retarget the integration away from enhancer and promoters but not enough to prevent the development of lymphomas by insertional mutagenesis. Our analysis points to the additional role of the enhancer/promoter in the MLV LTR to drive oncogenesis. Second generation SIN MLV vectors do take this feature into account and are associated with an increased safety profile (47). An *in vivo* analysis of BET-independent SIN MLV vectors is warranted. Such MLV vectors may have an increased genotoxic safety profile.

### Evolutionary advantage of the use of BET proteins for MLV integration

If the interaction of MLV IN with BET is dispensable for MLV replication *in vivo* and for lymphomagenesis, why did MLV evolve this mechanism? Although different explanations can be thought of, BET-mediated integration may provide the more efficient pathway. First, in the present study and the study published by the group of M. Roth (51), the number of integrations detected in mice infected with either W390A MLV or tail-peptide deleted MLV, was lower in comparison with the number of integrations for WT MLV. In this sense, BET-dependent MLV may be able to establish a higher number of proviruses inside the host genome, and a higher chance for a productive infection. Second, we show how BET proteins direct MLV integration *in vivo* towards enhancers and promoters. Integration at these sites ensures a high transcriptional activity. In fact, BRD4 can remain bound to several loci to remember the gene transcriptional profile after mitosis (78), a process MLV relies on for completing nuclear entry. Integration at these sites may guarantee MLV that the transcriptional machinery can initiate the transcription of the MLV genome and continue the replication cycle. The strong enhancer/promoter of the MLV LTR clearly contributes to the high transcriptional activity which is, in contrast to HIV, independent from a trans-activator such as Tat. Lymphomagenesis may not be an intrinsic feature of the MLV replication cycle but a bystander effect. Although this is not beneficial for a pathogen, in the case of mice the horizontal transmission to offspring is guaranteed before their death considering how early in their lifespan mice reach adulthood. The integration preference near enhancers and promoters and the strong LTR promoter all contribute to the risk for lymphomagenesis. Whether integration near oncogenes is stimulated by BRD4 interaction or BRD4-mediated integration near oncogenes is selected for clonal proliferation is not fully clear. Our data points towards the second scenario, but more detailed analyses are required. The multi-pronged mechanism to ensure proviral expression after transcription and thus the relative absence of latency may be associated with the intrinsic risk for insertional mutagenesis, clonal expansion and tumorigenesis.

## ACKNOWLEDGEMENTS

This work was supported by the Research Foundation Flanders (FWO) (G0A5316N and SBO-SAPHIR) and the KU Leuven Research Council (C14/17/095-3M170311). DVL, SVB, AB and SEA all received doctoral fellowships of the FWO. FC is an Industrial Research Fund (IOF) fellow. We would like to thank Dr. Alexandar Tzankov, for preparation of histology samples of lung, liver and kidney; Prof. Nancy Boeckx (KU Leuven) for the analysis of blood smears, as well as Dr. Susan R. Ross, for critical review and providing us the p63.2 MLV construct. We would like to thank Servier Medical Art for allowing the free use of their pictures in scientific publications under the Creative Commons License. We thank Dr. S. Ross for critical reading.

## AUTHOR CONTRIBUTIONS

JDR, IN and ZD conceptualized the experiments of this manuscript; MM, DVL, SVB, and SEA performed the experiments and analyzed the data; IN and AB pooled and analyzed genetic data, LM, TT and JS performed the histology analysis; RG, FC and ZD supervised experimental work. IN wrote the first draft of the manuscript; ZD supervised the project and reviewed the final version of the manuscript. All authors read and approved the final manuscript.

---

**Supplementary Figure 1.**
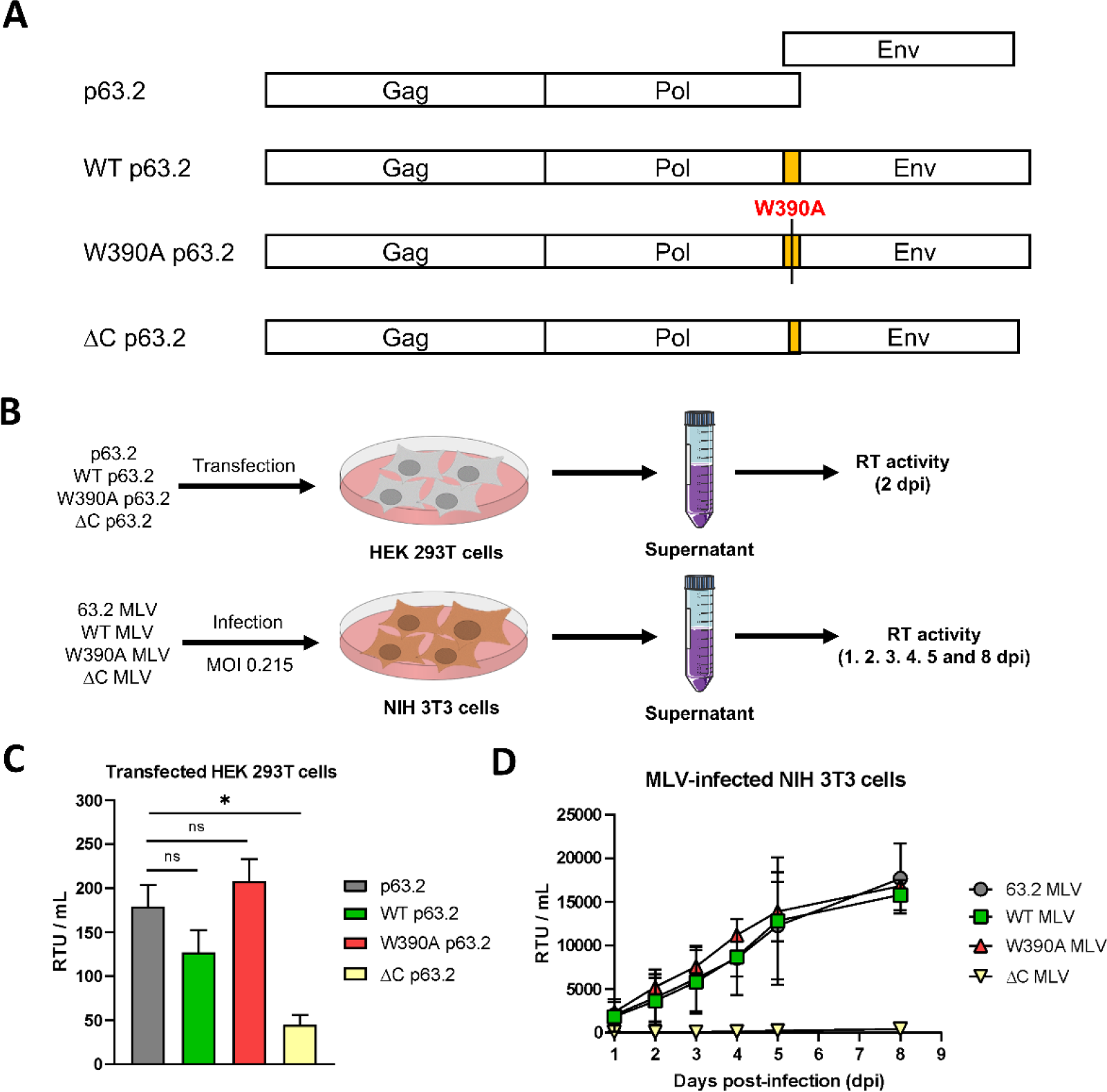
*In vitro* replication of MLV molecular clones. (**A**) Schematic representation of the p63.2, WT p63.2, W390A p63.2 and ΔC p63.2 molecular clones. p63.2 represents the wild type virus showing the overlap of *Pol* and *Env* genes (56). A codon optimized sequence duplicating the overlapping sequence at the protein level was inserted in front of the *Env* start codon (represented in orange) in molecular clone WT p63.2. In an identical way, the integrase W390A mutation was introduced in W390A p63.2. A fourth molecular clone was created containing a 27 amino acid IN C-terminal truncation (ΔC p63.2). (**B**) Schematic representation of the experiments to evaluate the replication of MLV molecular clones in HEK 2293T and NIH 3T3 cells. (**C**) Reverse transcriptase activity units (RTU) in the supernatants of HEK 293T cells transfected with p63.2, WT p63.2, W390A p63.2 or ΔC p63.2 molecular clones after two days post-transfection (n = 2). Statistical significance was analyzed using an ANOVA test. (**D**) Viral breakthrough in NIH 3T3 cells infected with 63.2 MLV, WT MLV, W390A MLV and ΔC MLV at MOI 0.215. Supernatants were collected after 1, 2, 3, 4, 5 and 8 days post- infection and RT activity was measured (n = 2).

**Supplementary Figure 2.**
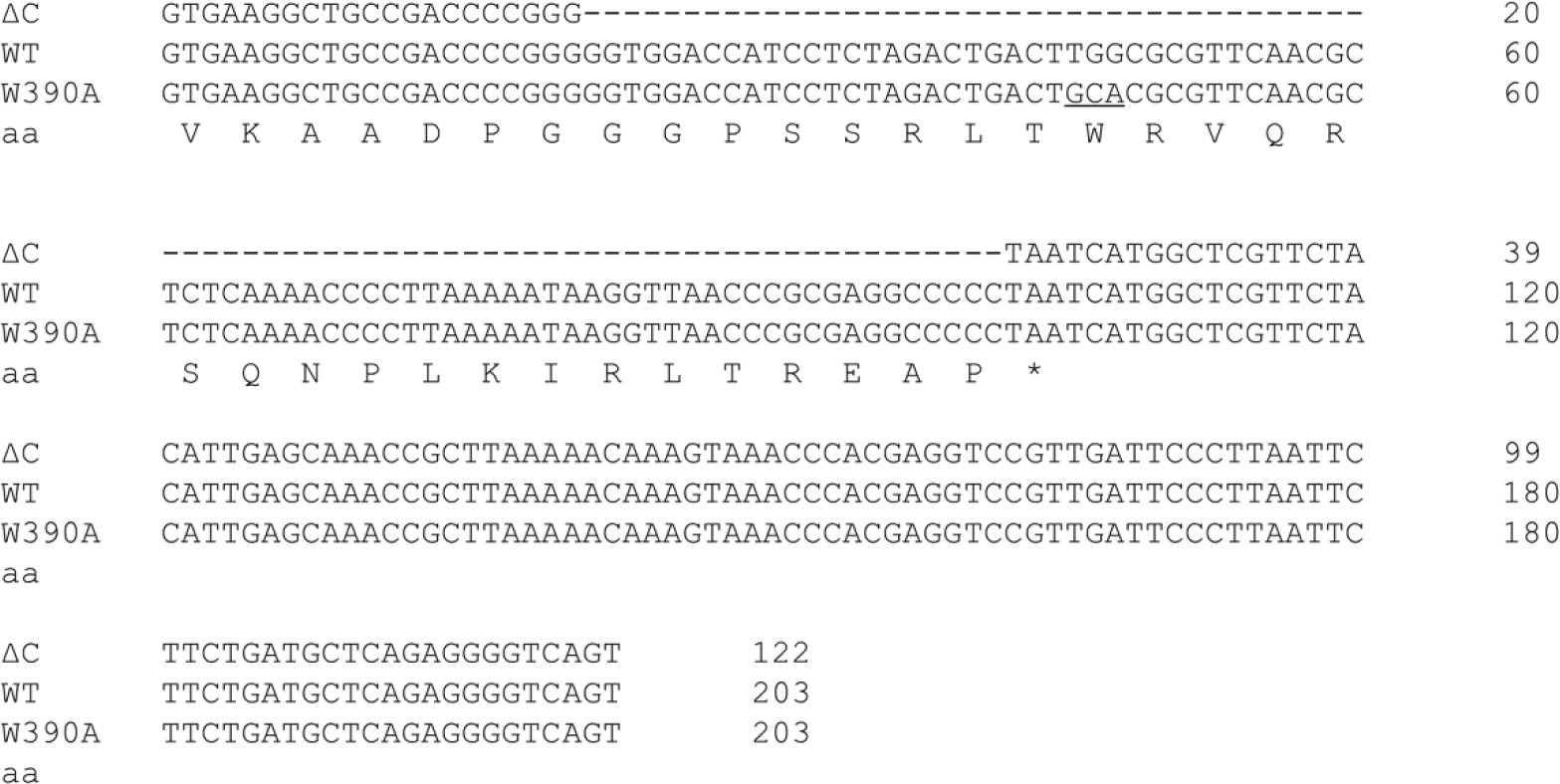
C-terminal integrase sequences of the ΔC p63, WT p63.2 and W390A p63.2 molecular clones. The W390A mutation is underlined. Amino acids (aa) coded by each codon are shown below nucleotides sequences. The stop codon is depicted with an asterisk (*). Deletions of nucleotides are indicated with a dash (–).

**Supplementary Figure 3.**
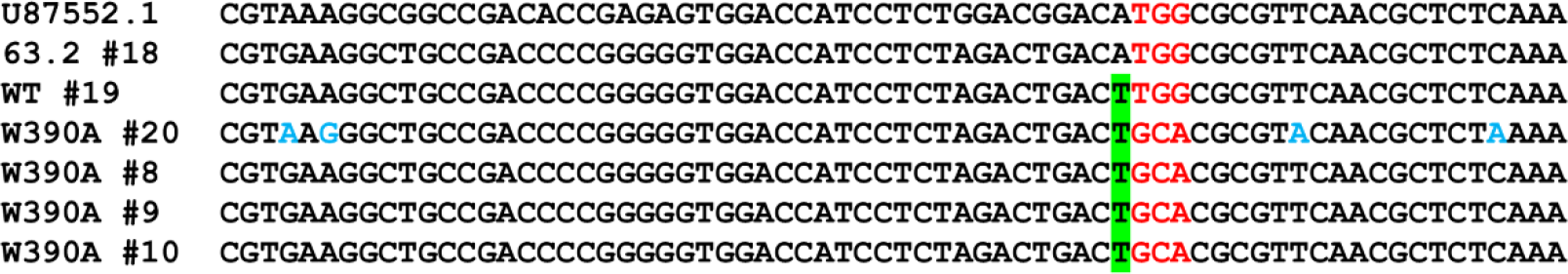
Revertant analysis performed by genomic DNA sequencing of six mice infected either with 63.2 MLV (n=1), WT MLV (n=1) or W390A MLV (n=4). U87552.1 indicates the reference MLV *Pol* gene sequence. Mice labelled as #18, #19 and #20 correspond to gDNA extracted from spleen of a mouse infected with 63.2 MLV, WT MLV or W390A after 12 weeks post-infection. Mice labeled #8, #9 and #10 were infected with W390A MLV and gDNA from spleen was extracted between 12-14 weeks post-infection. Nucleotides related to the MLV IN position 390 are colored in red. Nucleotides highlighted in green indicate a synonymous mutation observed in mice infected with WT MLV and W390A MLV near the 390 position. Other mutations in mice #20 are highlighter in blue.

**Supplementary Figure 4.**
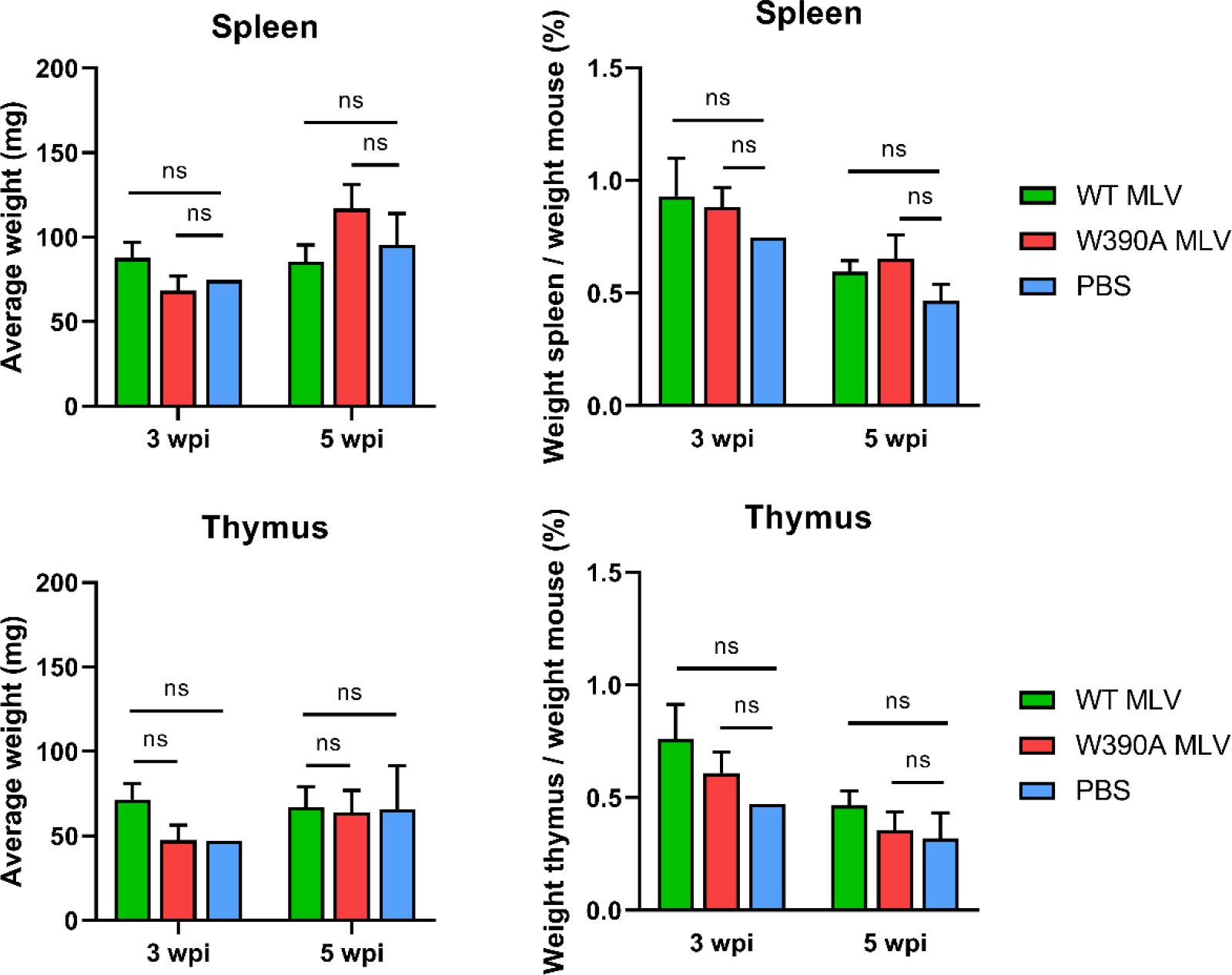
Average weight of the thymus and spleen from mice infected with 4·10^6^ RTU of WT MLV (n = 3) or W390A MLV (n = 3) after 3 and 5 weeks post- infection. Mice injected with PBS (n = 3) were used as negative controls. Bars show mean ± SD. No statistically significant differences were found using a Kluskal-Wallis test.

**Supplementary Figure 5.**
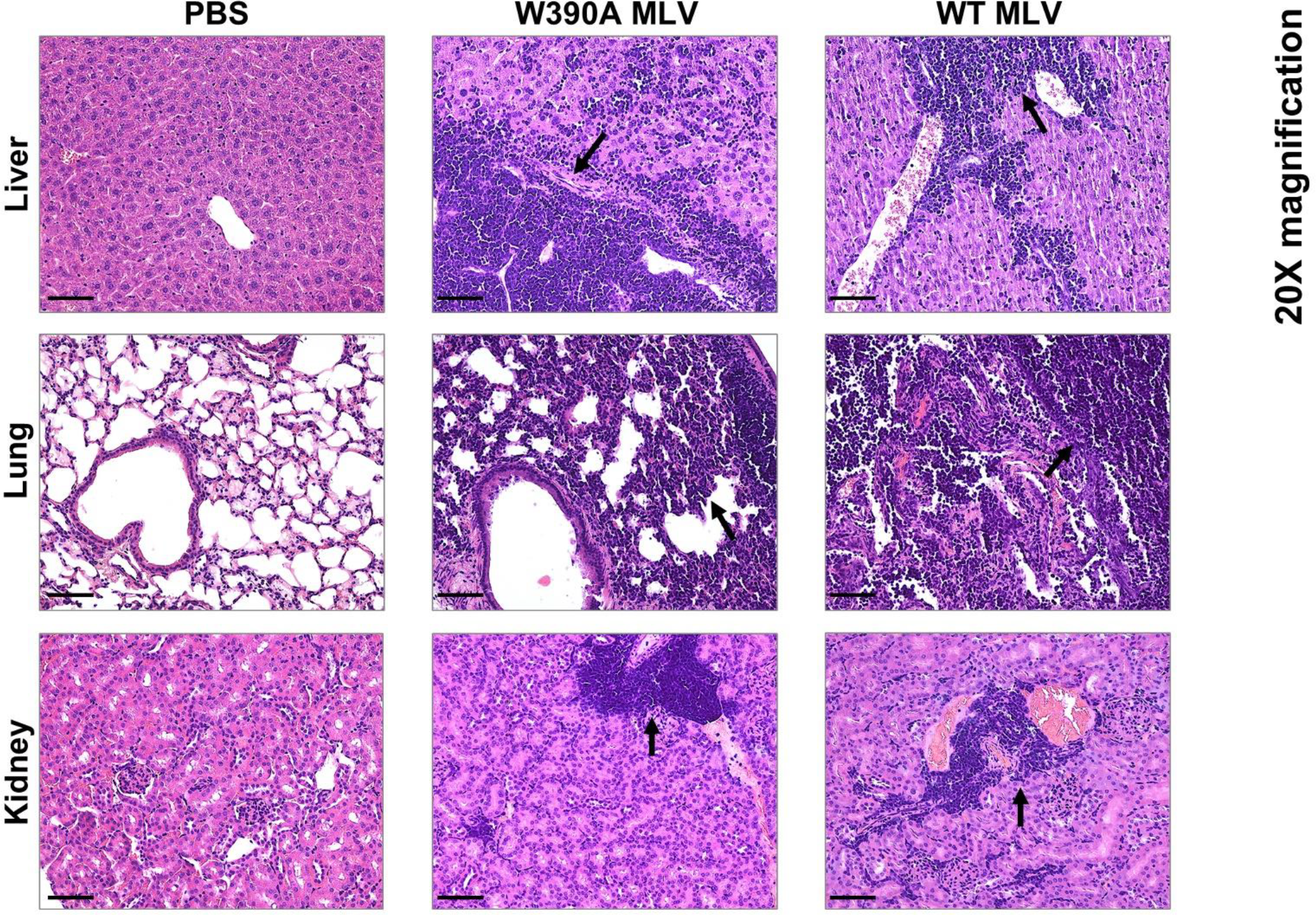
Hematoxylin and eosin staining of lung, kidney and liver tissues from mice infected with WT MLV or W390A MLV. Mice injected with PBS were used as negative control. Black arrows point towards zones with tumor infiltration. Scale bar indicates 40 µm.

**Supplementary Figure 6.**
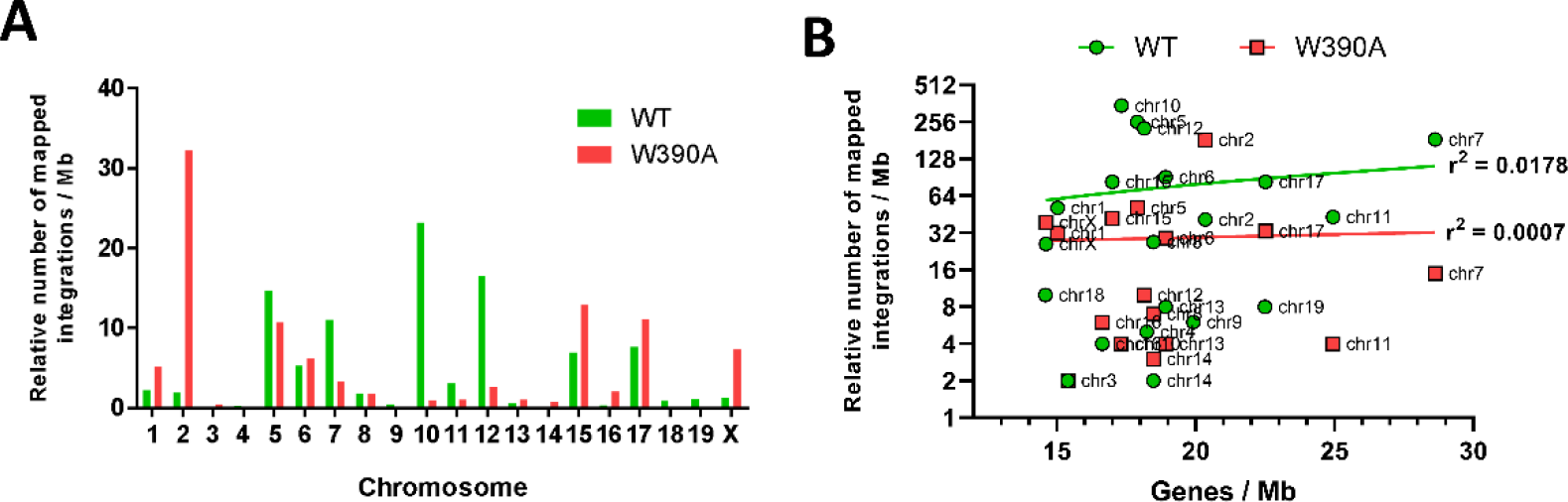
(**A**) Chromosome analysis summarizing mapped integration sites from WT MLV and W390A MLV into each mouse chromosome. (**B**) Relation between gene density and relative number of mapped integrations of WT MLV and W390A MLV. Each dot represents a single chromosome. Linear regression was used to analyze the relation between gene density and the number of mapped integrations. Mapped integrations for each chromosome were normalized to its corresponding size (megabase, Mb).

**Supplementary Figure 7.**
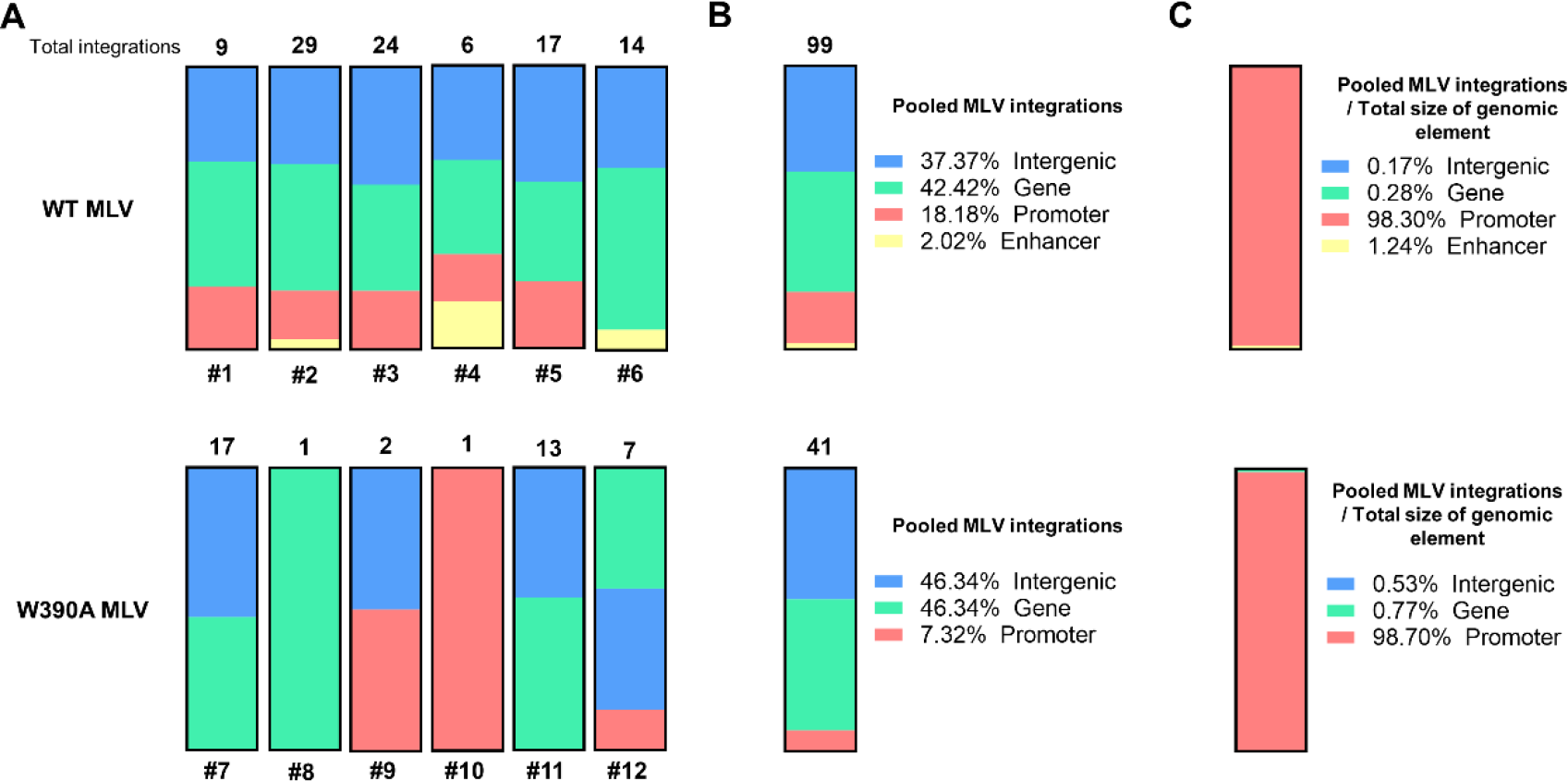
Integration site preference in clones generated by an infection with a lower dose of WT MLV or W390A MLV at late stage in the infection. The abundance of clones from mouse #1 to #12 was normalized to 1 to eliminate the expansion potential of MLV integration. (A) Integration preference into intergenic positions (blue), gene bodies (green), promoters (red) and enhancers (yellow) in WT MLV- and W390A MLV-infected mice. The numeric mouse label is indicated below each bar. Total number of integration events from clonal integration sites in each mouse is indicated on top of the bar. (B) Pooled integration preference into intergenic positions, gene bodies, promoters and enhancers in WT MLV- (n = 6) and W390A MLV-infected mice (n = 6). Total number of integrations events is indicated on top of the bar. (C) Integration preference into intergenic positions, gene bodies, promoters and enhancers normalized by the total genomic size of each genomic feature. Data was normalized by dividing the percentage of integration site preference into each genomic feature by its own estimated total genomic size: intergenic positions (58.97%), genes (40.54%), promoters (0.05%) and enhancers (0.44%).

**Supplementary Figure 8.**
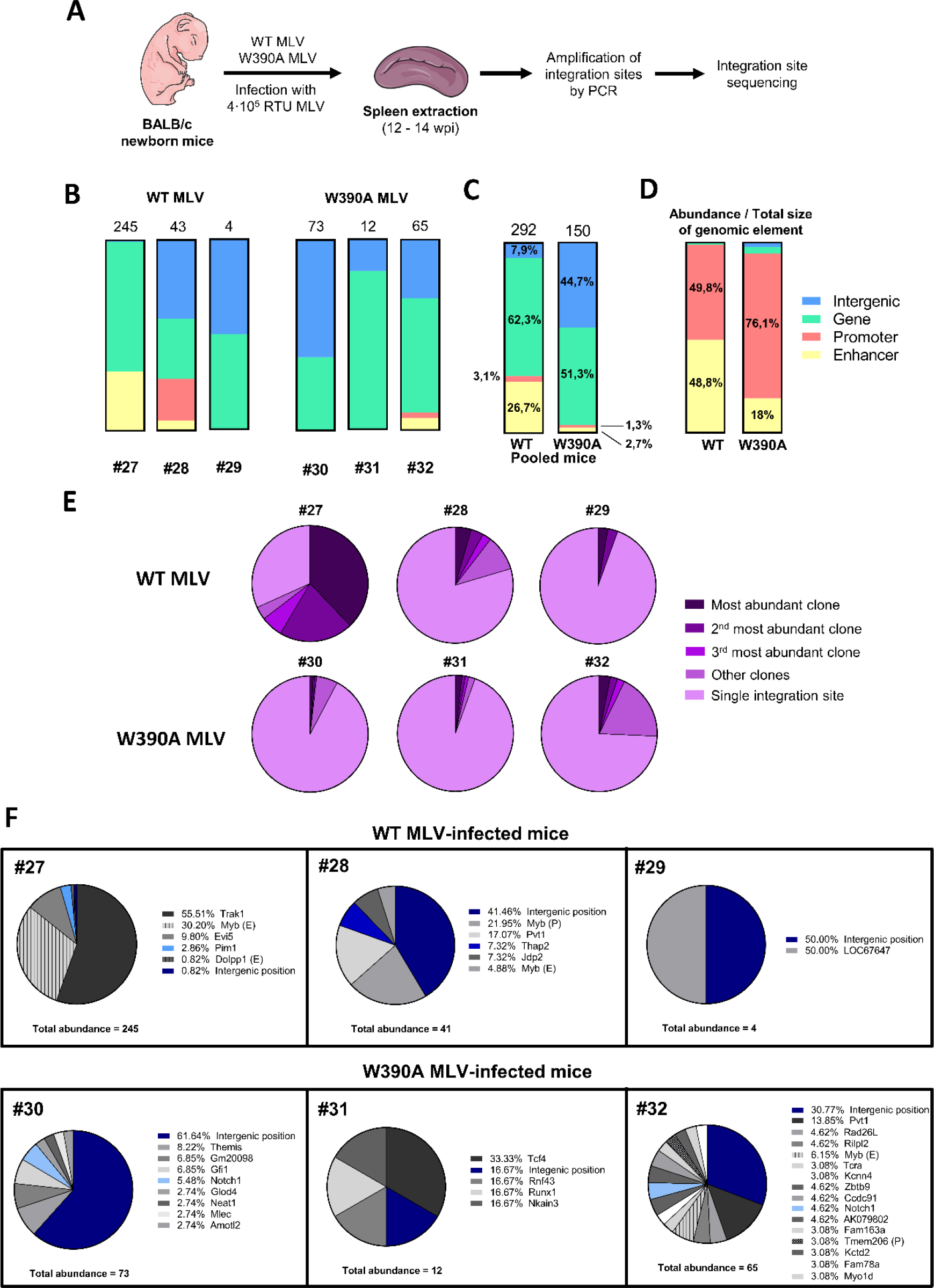
Distribution of MLV abundance in clones generated by an infection with a lower dose of WT MLV or W390A MLV. (**A**) Schematic representation of the work flow to determine the integration sites and the epigenetic features of WT MLV and W390A MLV integrants *in vivo* (**B**) MLV abundance into intergenic positions (blue), gene bodies (green), promoters (red) and enhancers (yellow) in WT MLV- and W390A MLV-infected mice. The numeric mouse label is indicated below each bar. Total abundance of clonal integration sites in each mouse is indicated on top of the bar. (**C**) Pooled MLV abundance in intergenic positions (blue), gene bodies (green), promoters (red) and enhancers (yellow) in WT MLV- (n = 3) and W390A MLV-infected mice (n = 3). Total abundance of pooled clonal integration sites from mice infected with either WT MLV or W390A MLV at a lower dose is indicated on top of the bar. Distribution of integration sites between WT MLV and W390A MLV was analyzed using a chi-squared test. The test reported a statistically significant difference between both distribution (*P*-value ≤ 0.0001). (**D**) Total abundance from intergenic positions (blue), gene bodies (green), promoters (red) and enhancers (yellow) normalized by the total genomic size of each genomic feature. Data were normalized by dividing the percentage of integration sites into each genomic feature by its own estimated total genomic size: intergenic positions (58.97%), genes (40.54%), promoters (0.05%) and enhancers (0.44%). Data represent six mice infected with in WT MLV or W390A MLV. (**E**)Pie charts representing the relative abundance of clones and single integration sites. Each pie chart corresponds to a single mouse infected with WT MLV or W390 MLV. (**F**) Pie charts depicting individual gene insertional profile of splenic cells with clonal expansion from mice infected with 4⋅10^5^ RTU of either WT MLV or W390A MLV. Total abundance detected by NGS analysis is shown below each pie chart for both MLV constructs. The percentage before each gene name indicates the relative abundance of this gene. Genes with insertions of both WT MLV and W390A MLV are colored in a blue scale. Insertions into enhancers or promoters are indicated with an E or P after gene name, respectively.

**Supplementary Figure 9.**
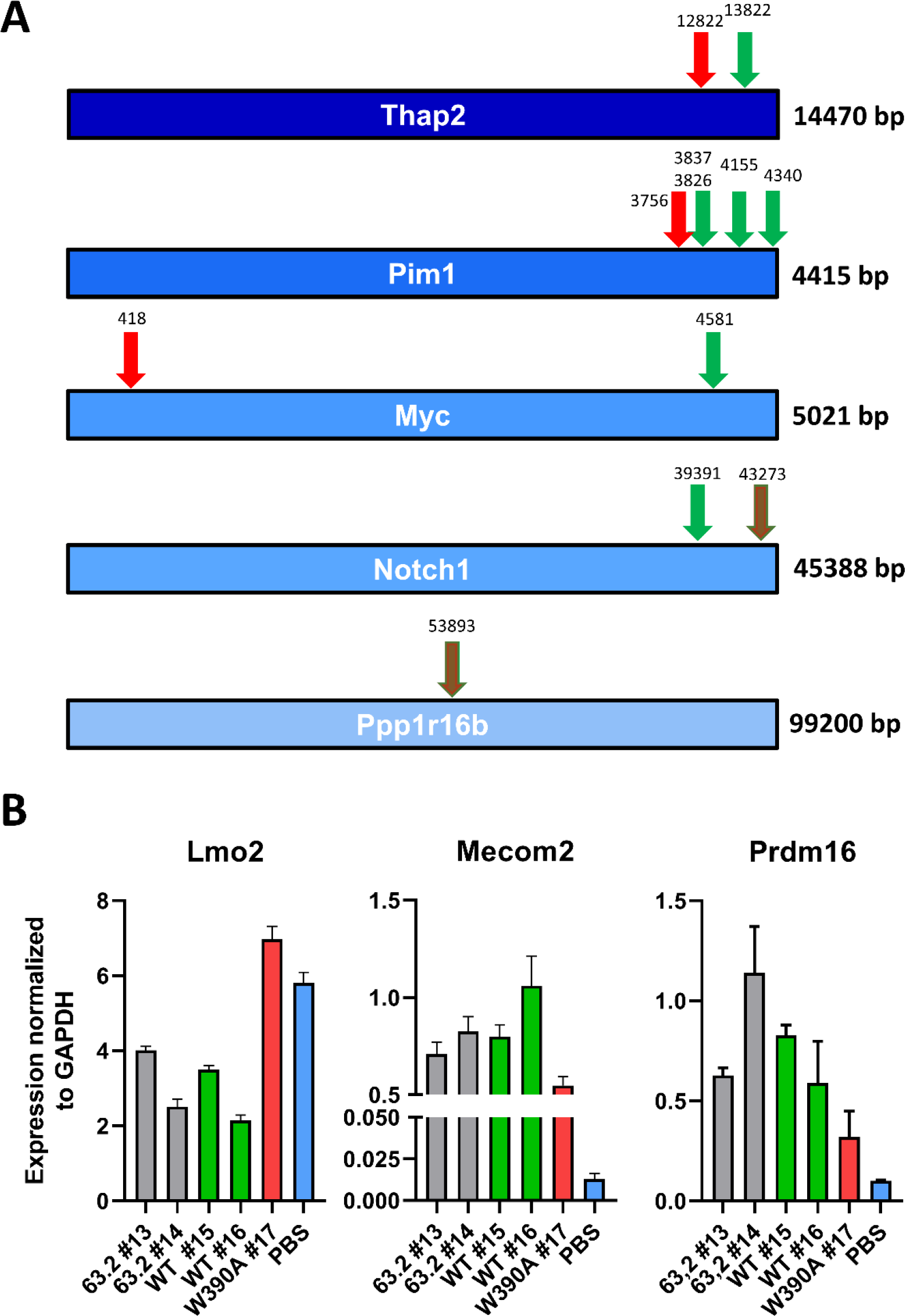
Mechanisms of insertional mutagenesis. (**A**) WT and W390A MLV insertions in oncogenes. WT MLV and W390A MLV insertional profile into *Thap2*, *Pim1*, *Myc*, *Ppp1r16b* and *Notch1* genes. Green arrows indicate WT MLV insertions while red arrows indicate W390A MLV insertions. Numbers above each arrow indicate the base pair position of the integration. Brown arrows indicate insertion of both WT MLV and W390A MLV. (**B**) MLV-LTR driven expression of oncogenes. RT-qPCR analysis of expression of Lmo2, Mecom2 and Prdm16 genes in spleen cells from mice infected with 63.2 MLV, WT MLV or W390A MLV after 12-14 weeks post-infection. GAPDH was used as housekeeping gene. The PBS condition was used as a negative control.

